# Singlet oxygen-dependent chloroplast degradation is independent of macroautophagy in the *Arabidopsis ferrochelatase* two mutant

**DOI:** 10.1101/2021.02.17.431731

**Authors:** Matthew D. Lemke, Karen E. Fisher, Marta A. Kozlowska, David Tano, Jesse D. Woodson

## Abstract

**Background:** Chloroplasts respond to stress and changes in the environment by producing reactive oxygen species (ROS) that have specific signaling abilities. The ROS singlet oxygen (^1^O_2_) is unique in that it can signal to initiate selective degradation of damaged chloroplasts and then cell death. This chloroplast quality control pathway can be monitored in the *Arabidopsis* mutant *plastid ferrochelatase two* (*fc2*) that conditionally accumulates chloroplast ^1^O_2_ under diurnal light cycling conditions leading to rapid chloroplast degradation and eventual cell death. The cellular machinery involved in such degradation, however, remains unknown. Recently it has been demonstrated that whole damaged chloroplasts can be transported to the central vacuole via a process requiring autophagosomes and core components of the autophagy machinery. The relationship between this process, referred to as chlorophagy, and the degradation of ^1^O_2_-stressed chloroplasts and cells has remained unexplored.

**Results:** To further understand ^1^O_2_-induced cellular degradation and determine what role autophagy may play, the expression of autophagy-related genes were monitored in ^1^O_2_-stressed *fc2* seedlings and found to be induced. Although autophagosomes were present in *fc2* cells, they did not associate with chloroplasts during ^1^O_2_ stress. Mutations blocking the core autophagy machinery (*atg5, atg7*, and *atg10*) were unable to suppress ^1^O_2_-induced chloroplast degradation or cell death in the *fc2* mutant, suggesting autophagosome formation and macroautophagy are dispensable for ^1^O_2_–mediated cellular degradation. However, both *atg5* and *atg7* led to specific defects in chloroplast ultrastructure and photosynthetic efficiencies, suggesting macroautophagy may be involved in protecting chloroplasts from photo-oxidative damage. Finally, genes predicted to be involved in microautophagy were shown to be induced in stressed *fc2* seedlings, indicating a possible role for an alternate form of autophagy in the dismantling of ^1^O_2_-damaged chloroplasts.

**Conclusions:** Our results support the hypothesis that ^1^O_2_-dependent chloroplast degradation is independent from autophagosome formation, canonical macroautophagy, and chlorophagy. Instead, ATG-independent microautophagy may be involved in such degradation. However, canonical macroautophagy may still play a role in protecting chloroplasts from ^1^O_2_-induced photo-oxidative stress. Together, this suggests chloroplast function and degradation is a complex process that utilizes multiple autophagy and degradation machineries, possibly depending on the type of stress or damage incurred.

## Background

Plants have evolved intricate signaling systems that enable them to sense and subsequently respond to environmental stresses. One way plants achieve this is through the use of their chloroplasts (specialized photosynthetic plastid organelles) as sensors for multiple types of abiotic and biotic stresses. Even under ideal conditions, the unique photochemistry in chloroplasts leads to the production of the reactive oxygen species (ROS) superoxide and singlet oxygen (^1^O_2_) at photosystems I (PSI) and II (PSII), respectively (1). Plants have evolved mechanisms to combat this ROS production, including non-photochemical quenching (NPQ), the production of ROS quenching pigments and enzymes, and systems to regulate energy capture and photosynthetic activity (2). However, chloroplast ROS may further accumulate under various stresses including excess light (EL) (3), drought (4), salinity (5), and pathogen attack (6). In such cases, this ROS may then overwhelm the protective mechanisms and lead to chloroplast and photosystem damage though the oxidation of proteins, DNA, and lipids (3). At the same time, chloroplast ROS also act as signaling molecules (4,7). Both hydrogen peroxide (produced after the dismutation of superoxide) and ^1^O_2_ have been demonstrated to induce separate signals to control the expression of hundreds of nuclear genes (8). Moreover, ^1^O_2_ has been shown to induce programmed cellular degradation, including selective chloroplast degradation (9,10).

The ability of ^1^O_2_ to induce such cellular degradation has been inferred from multiple *Arabidopsis* mutants that specifically and conditionally accumulate large amounts of chloroplast ^1^O_2_ (11). One such mutant, *plastid ferrochelatase two* (*fc2*), is defective in one of the two conserved chloroplast enzymes required for heme synthesis. Under diurnal light cycling conditions, this mutant accumulates the substrate of the FC2 enzyme, protoporphyrin IX (PPIX), an intermediate of the tetrapyrrole (e.g., chlorophylls and hemes) biosynthetic pathway (12). Like other unbound tetrapyrroles, the accumulation of free PPIX leads to the rapid production of ^1^O_2_ in the light (13). Within hours, such high ^1^O_2_ levels lead to wholesale chloroplast degradation followed by cell death. This can be easily visualized in seedlings or adult plants by the rapid bleaching of cotyledons and necrotic lesions in leaves, respectively (14,15). Even when *fc2* mutants are grown under permissive 24h constant light conditions that avoid cell death, chloroplasts are still observed to degrade within the cytoplasm. These degrading chloroplasts can sometimes be observed protruding or blebbing into the central vacuole of an otherwise healthy cell. Together, it was hypothesized that certain chloroplasts are selectively degraded in this mutant (14,15). The mechanisms controlling this degradation and the subsequent transport of chloroplast material into the central vacuole, however, are unknown.

^1^O_2_ is naturally produced at PSII during EL and other environmental stresses that inhibit photosynthesis (3,16). Under such conditions, chlorophylls in PSII become unable to transfer their energy to the photosynthetic reaction centers, excite to their triplet state, and interact with molecular oxygen to generate ^1^O_2_. Unlike hydrogen peroxide that is simultaneously made during most of these stresses, ^1^O_2_ has an extremely short half-life (0.5-1 μsec in cells (17)) and is likely confined to the same chloroplast in which it was generated (^1^O_2_ can travel < 200 nm, but chloroplasts are 5-10 μm wide). Therefore, secondary signals that control nuclear gene expression and/or chloroplast degradation in response to ^1^O_2_ accumulation likely exist. To identify such signaling factors involved in ^1^O_2_-induced chloroplast degradation, a genetic screen for *ferrochelatase two suppressor* (*fts*) mutations that block cell death and chloroplast degradation in the *fc2* mutant have been isolated (14). Two *fts* mutants have been shown to affect *PENTATRICOPEPTIDE-REPEAT-CONTAINING PROTEIN 30* (*PPR30*) and “*mitochondrial*” *TRANSCRIPTION TERMINATION FACTOR 9* (*mTERF9*) (15). Both genes encode chloroplast-localized proteins predicted to be involved in post-transcriptional gene regulation; PPR30 belongs to the P-class of PPR proteins that regulate gene expression by directly affecting RNA transcript stability, processing, editing, and/or translation (18) and mTERF9 has been demonstrated to be required for assembly of the 30S ribosome subunit in chloroplasts (19). As such, these results suggest a (currently unidentified) chloroplast-encoded factor(s) may be required to initiate the ^1^O_2_ signal.

A third *fts* mutant was shown to affect the cytoplasmic ubiquitin E3 ligase Plant U-Box 4 (PUB4) suggesting the cellular ubiquitination machinery is involved in targeting damaged chloroplasts for degradation (14). Consistent with this hypothesis, proteins associated with ^1^O_2_-stressed chloroplasts became ubiquitinated prior to degradation. This degradation was mostly blocked in the *fc2 pub4* double mutant, suggesting PUB4, or another E3 ligase may mark damaged chloroplasts for turnover by ubiquitination (14). As such, chloroplast generated ^1^O_2_ may induce a chloroplast quality control system allowing cells to sense and turnover damaged chloroplasts to sustain healthy chloroplast populations and maintain efficient energy production. Which cellular machinery may be involved in the targeted degradation and removal of these chloroplasts, however, remains unknown.

Cellular quality control systems are fundamental mechanisms in biology that allow organisms to maintain the optimal functionality of molecular processes. In the case of damaged proteins, cellular components, or even whole organelles, autophagy often plays a role in such processes (20,21). A conserved process in eukaryotes, autophagy allows cells to degrade and recycle macromolecules and larger cellular components such as whole organelles (22,23). In yeast and mammals, ROS-induced mitochondrial degradation (mitophagy) is regulated by the E3 ubiquitin ligase PARKIN, leading to the autophagic removal of mitochondria to the lysosome (24).

In eukaryotic cells, two major types of autophagy are used; macroautophagy and microautophagy. Macroautophagy involves the selective sequestration of proteins and organelles by packaging such cytosolic components into autophagosomes, which are then transported to the central vacuole for turnover (25,26). Macroautophagy is a well-studied and fundamental process that is strongly conserved across eukaryotic kingdoms (27). In plants, canonical macroautophagy is generally characterized as a process whereby cytosolic components are encapsulated in a double membrane vesicle, known as an autophagosome, and are then transported to the central vacuole for turnover and eventual remobilization of nutrients. This process is fundamental in the maintenance of cellular homeostasis, nutrient storage, and degradation of cytotoxic components and dysfunctional proteins (28,29). This form of autophagy has been well characterized to proceed in an Autophagy 5- (ATG5) and Autophagy 7- (ATG7) dependent manner, requiring core autophagy proteins to play key roles in the Autophagy 8 (ATG8) conjugation system, a process that drives the formation and maturation of autophagic bodies which encapsulate and package cytosolic components (30). Alternatively, cytosolic components can be removed by invagination of the vacuole in processes resembling endocytosis via microautophagy (31,32). Compared to macroautophagy, significantly less is known about this process, particularly in plants (31).

Recent work has suggested whole chloroplasts are also transported to the central vacuole for degradation, either by processes that resemble macroautophagy or microautophagy, after damage by UVB (33) or excess light (EL) (34), respectively. Under UVB stress, damaged chloroplasts are completely engulfed by autophagosomes, a hallmark of macroautophagy. Under EL stress, however, only partial autophagosomes associate with damaged chloroplasts. This process resembles microautophagy, where cytoplasmic components are transported into the central vacuole without complete envelopment by an autophagosome (34). The specific mechanisms involved in these processes (collectively dubbed chlorophagy) remain unclear, but it is evident that both are dependent on core autophagy proteins. Under both UVB and EL stress, the transport of damaged chloroplasts to the central vacuole is blocked in *atg5* and *atg7* mutants, suggesting that autophagic bodies are required to proceed and complete chlorophagy (34,35). Recent work has shown that PUB4 is dispensable to chlorophagy during EL stress, suggesting that this E3 ligase is not necessary for autophagosome formation (36). However, the relationship between autophagy and chloroplast quality control during ^1^O_2_ stress and has remained unexplored.

In this study, we sought to further understand the mechanisms involved in ^1^O_2_-induced chloroplast quality and cell death in *fc2* mutants by determining what role autophagy may play in these pathways. Although a large number of autophagy-related genes are induced in stressed *fc2* seedlings, autophagosomes do not interact with these ^1^O_2_-stressed chloroplasts. Moreover, mutations affecting the core autophagy machinery do not block chloroplast turnover or cell death in the *fc2* mutant. Finally, our gene expression analyses point towards a potential role for microautophagy in ^1^O_2_-induced chloroplast quality control. Together, these results suggest that multiple chloroplast degradation pathways likely exist in plant systems and that chloroplast quality control itself is likely a complex process regulated by multiple types of cellular autophagy machinery.

## Results

### Autophagy is transcriptionally induced in stressed *fc2* mutant seedlings

In *fc2* mutants, the accumulation of ^1^O_2_ in chloroplasts leads to their selective degradation (14). In such cases, degrading chloroplasts appear to interact with the central vacuole where they may be further degraded and recycled. While UVB and EL damaged chloroplasts have been shown to be transported to the vacuole in an autophagy-dependent manner (34,35), the cellular machinery involved in transporting ^1^O_2_-damaged chloroplasts in the *fc2-1* mutant is unknown.

To investigate if autophagy may also be involved in degrading chloroplasts in *fc2* mutants, we first tested if core autophagy and autophagy-related genes are transcriptionally induced by ^1^O_2_ accumulation. Using a previously published microarray data set of wild type (wt) and *fc2-1* seedlings before and during de-etiolation (14), we analyzed the expression of a manually curated list of 71 core autophagy and autophagy-related genes (Fig. 1a and Table S1). Compared to wt, the majority (85 %) of these genes were induced in the *fc2-1* mutant (with no cutoff values applied) and the 120 minute time point showed the strongest induction (72% of genes). In this time point, the top five most highly induced autophagy-related genes were *BAG6* (*AT2G46240*), *WRKY33* (*AT2G38470*), *RAB7* (AT1G22740), *ATG1B* (*AT3G53930*), and *CDC48C* (*AT5G03340*) (Fig. 1b). Of particular interest is *BAG6*, a strongly conserved positive regulator of programmed cell death (37). To determine if *fc2-1* seedlings respond similarly under light cycling conditions, we monitored the expression of these five genes in four-day old seedlings grown in 6h light/18h dark cycling light conditions. Compared to wt, four of the five genes (excepting *RAB7*) were significantly induced in *fc2-1* seedlings suggesting these expression patterns are maintained post de-etiolation (Fig. 1c).

**Figure 1.**
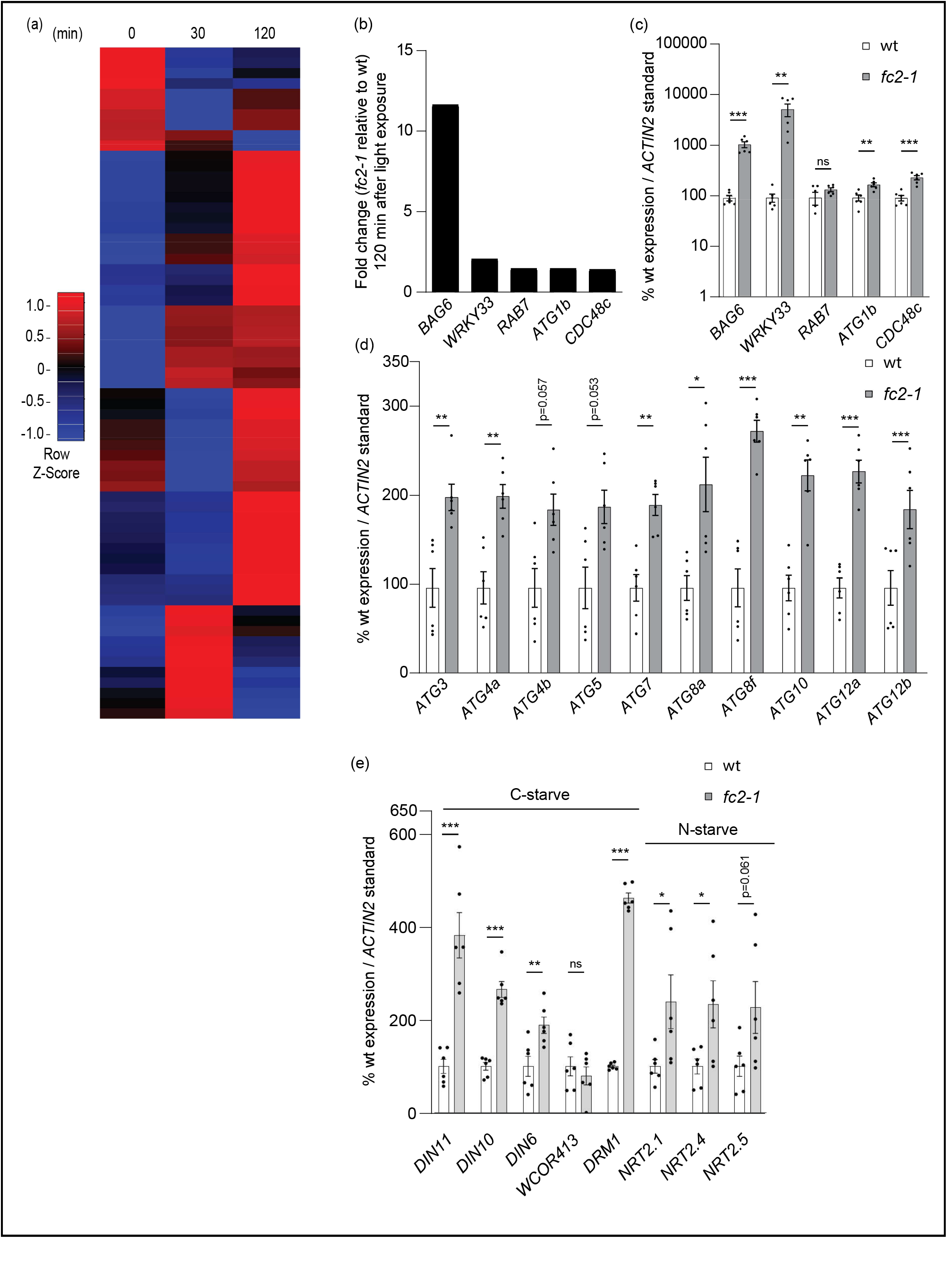
Autophagy-related genes are transcriptionally induced in stressed *fc2-1* seedlings. The expression of autophagy-related genes were monitored in *fc2-1* mutants. **A)** A heatmap of autophagy-related gene expression (relative to wt) in etiolated seedlings at time points prior to (0 min) and during de-etiolation (30 min and 120 min). Microarray data was previously generated from etiolated seedlings grown in the dark for four days and exposed to light for the indicated amount of time (14). The list of genes considered is presented in Table S1. **B)** Expression fold change (relative to wt) of the top five most up-regulated core autophagy and autophagy genes at 120 minutes from panel A. **C), D)**, and **E)** RT-qPCR analysis of autophagy-related, core autophagy, and starvation (carbon and nitrogen) marker transcripts, respectively, from four-day-old seedlings grown under 6h light/18h dark light cycling conditions. Shown are mean values +/- SEM (n = 6 biological replicates). Statistical analyses were performed by student’s t-tests. *, **, *** indicate a p-value of ≤ 0.05, ≤ 0.01, and ≤ 0.001, respectively. In all bar graphs, closed circles represent individual data points.

As autophagy-related genes appear to have a general induction pattern in stressed *fc2-1* seedlings, we probed ten core autophagy genes that make up the ATG8-conjugation system to explore if macroautophagy may be transcriptionally induced. Again, we saw a similar trend in the induction of these genes under 6h light/18h dark cycling light conditions. All ten genes (eight significantly, p values ≤ 0.05) were induced in *fc2-1* compared to wt (Fig. 1d). Together, these data demonstrate that a subset of autophagy-related genes, including those that compose the ATG8-conjugation system, are transcriptionally induced in ^1^O_2_-stressed *fc2-1* seedlings during and after de-etiolation and suggest macroautophagy may be activated in response to chloroplast ^1^O_2_ stress.

As photosynthesis is likely impaired in *fc2-1* seedlings (12,14) and as carbon (C) and nitrogen (N) starvation have been shown to play a role in the induction of autophagy (38) we explored the possibility that *fc2-1* seedlings are exhibiting a starvation response. To this end, we probed a selection of C- and N-starvation marker genes (38) in four day old seedlings grown under 6h light/18h dark cycling light conditions. Compared to wt, the C-starvation marker genes (*DIN11* (*AT3G49620*), *DIN10* (*AT5G20250*) (39), *DIN6* (*AT3G47340*), and *DRM1* (*AT1G28330*) (40)) were significantly induced in *fc2-1* seedlings (Fig. 1e). The expression of one marker gene, *WCOR413* (*AT4G37220*), was not significantly different between wt and *fc2-1*. To investigate an N-starvation response, we monitored three markers genes; *AT1G08090* (*NRT2*.*1*), *AT5G60770* (*NRT2*.*4*), *AT1G12940* (*NRT2*.*5*) (41). Two of these genes, *NRT2*.*1* and *NRT2*.*4*, were significantly induced in *fc2-1* compared to wt (Fig. 1e). Together, these results suggest that stressed *fc2-1* seedlings are experiencing C- and N-starvation responses that may contribute to the transcriptional induction of autophagy-related genes.

### ATG8 does not associate with stressed chloroplasts in *fc2* mutants

As autophagy-related genes are transcriptionally induced in *fc2-1* mutants, we next sought to determine if canonical macroautophagy itself is active in *fc2-1* seedlings and if autophagic bodies associate with ^1^O_2_ stressed chloroplasts. Such a response can be monitored by following the localization of ATG8 protein, a post-transcriptionally regulated hallmark of autophagosome formation and macroautophagy (42). During chlorophagy, for instance, GFP-ATG8a can be observed as punctae interacting with chloroplasts after UVB or EL stress (43,44).

To determine if ATG8 and autophagosomes also interact with ^1^O_2_ stressed chloroplasts in the *fc2-1* mutant, we introduced a *UBQ10*::*GFP-ATG8a* construct into the *fc2-1* background and monitored GFP punctae using confocal microscopy. As a control, we tested if GFP-ATG8a still associated with senescing chloroplasts during dark-induced carbon starvation as previously reported (45). Under both 24h constant and 6h light/18h dark cycling light conditions, GFP punctae were observed in *fc2* cells, but did not appear to interact with chloroplasts under either condition (Fig. 2a). Under dark-grown starvation conditions however, GFP-ATG8 did associate with chloroplasts (Fig. 2a). Together, these results suggest autophagosomes are present in *fc2-1* seedling mesophyll cells, but are not associating directly with ^1^O_2_ stressed chloroplasts.

**Figure 2.**
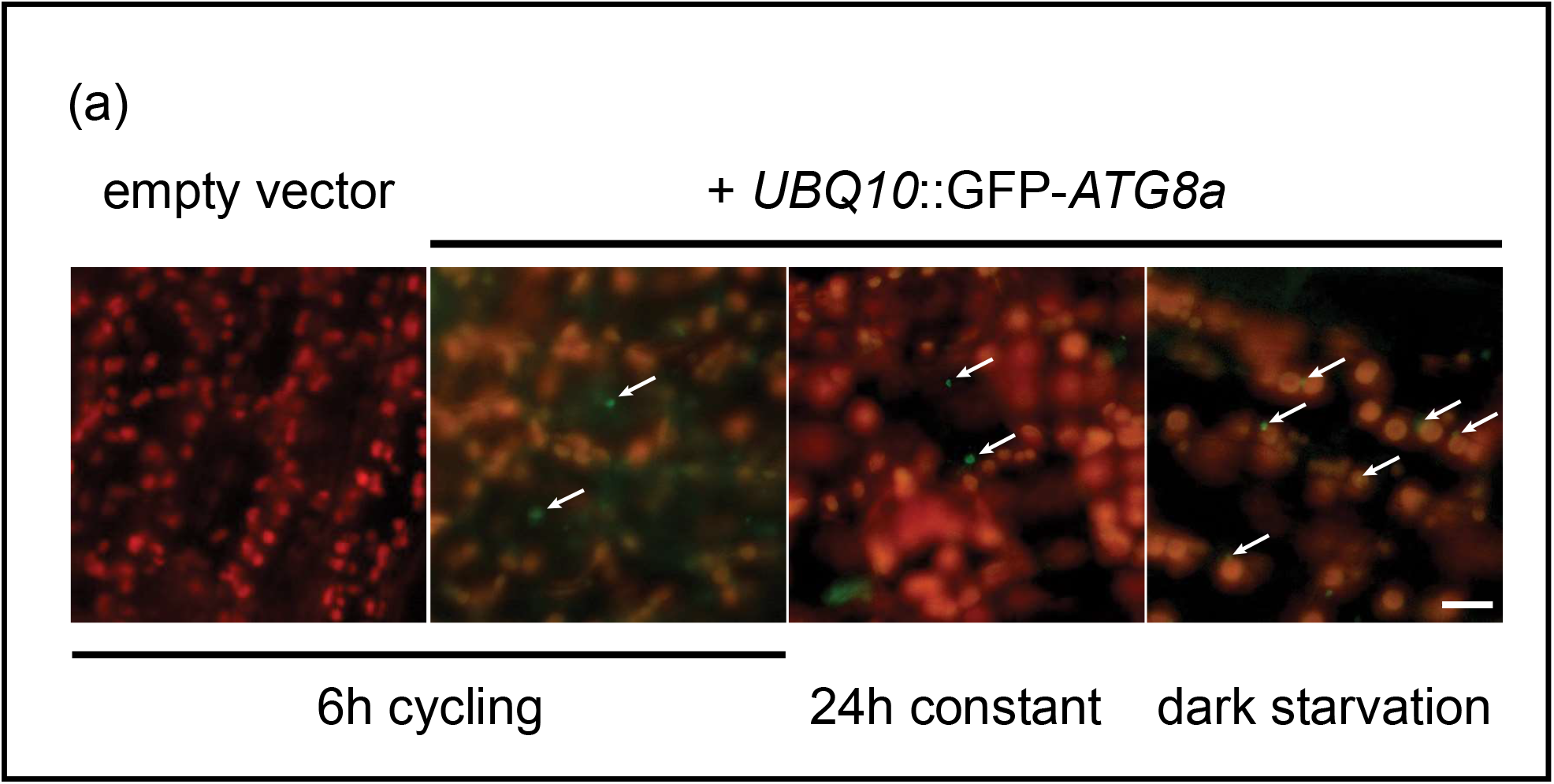
ATG8a does not associate with ^1^O_2_-stressed chloroplasts in *fc2* seedlings. The localization of GFP-ATG8a was assessed in stressed *fc2-1* seedlings. **A)** Shown are representative images of GFP-ATG8a in four day-old *fc2-1* seedlings grown in 24h constant light or 6h light/18h dark cycling light. The same line was also subjected to dark-induced carbon starvation for six additional days. Each panel is an overlay of GFP (green) and chlorophyll autofluorescence (red). White arrows indicate GFP-ATG8a punctae in cells. In the dark-starved sample, these punctae are associating with chloroplasts. Scale bars = 10 µm,

### Blocking macroautophagy does not suppress cell death in *fc2* seedlings

The above results suggest autophagosomes are active in *fc2* cells, but do not directly target chloroplasts for degradation. However, it is also possible that such interactions are too transient to detect, or that only a small portion of autophagosomes are involved in chloroplast stress. Therefore, to further investigate the role of macroautophagy in ^1^O_2_-mediated chloroplast degradation, we tested if there is a genetic interaction between chloroplast quality control and macroautophagy in *fc2* seedlings. In *Arabidopsis*, core autophagy (ATG) proteins such as ATG5, ATG7, and Autophagy 10 (ATG10), have been shown to be essential for autophagosome formation and thus are required for canonical macroautophagy (46–48). In addition, both ATG5 and ATG7 have been shown to be specifically required for chlorophagy (34,35). Therefore, we introduced null *atg5-1, atg7-2*, and *atg10-1* mutations into the *fc2-1* background. An RT-qPCR analysis confirmed the three *fc2-1 atg* double mutants lacked their respective *ATG* transcripts (Fig. S1a) and exhibited the expected early senescence phenotypes caused by these mutations (Fig. S1b) (46,48,49).

Next, we tested the ability of these *atg* mutations to suppress the *fc2-1* cell death phenotype, which can be achieved by reducing ^1^O_2_ accumulation (class I mutants such as *toc33*) or by blocking the ^1^O_2_ signal (class II mutants such as *pub4-6*) (14). As expected under 6h light/18h dark cycling light conditions, *fc2-1* mutants suffer from cell death and fail to green (Fig. 3a). This was suppressed by *toc33* and *pub4-6*, but not by any of the three *atg* mutations. This cell death was dependent on light cycling as these mutants appeared healthy under permissive 24h constant light conditions. Furthermore, in the wt background, all three *atg* single mutants appeared healthy suggesting that cell death was dependent on the *fc2-1* background (Fig. S2a). Cell death itself was confirmed through trypan blue staining of the seedlings (Figs. 3b and c and S2b and c) and there was no significant difference between *fc2-1* and the *fc2-1 atg* double mutants under light cycling conditions.

**Figure 3.**
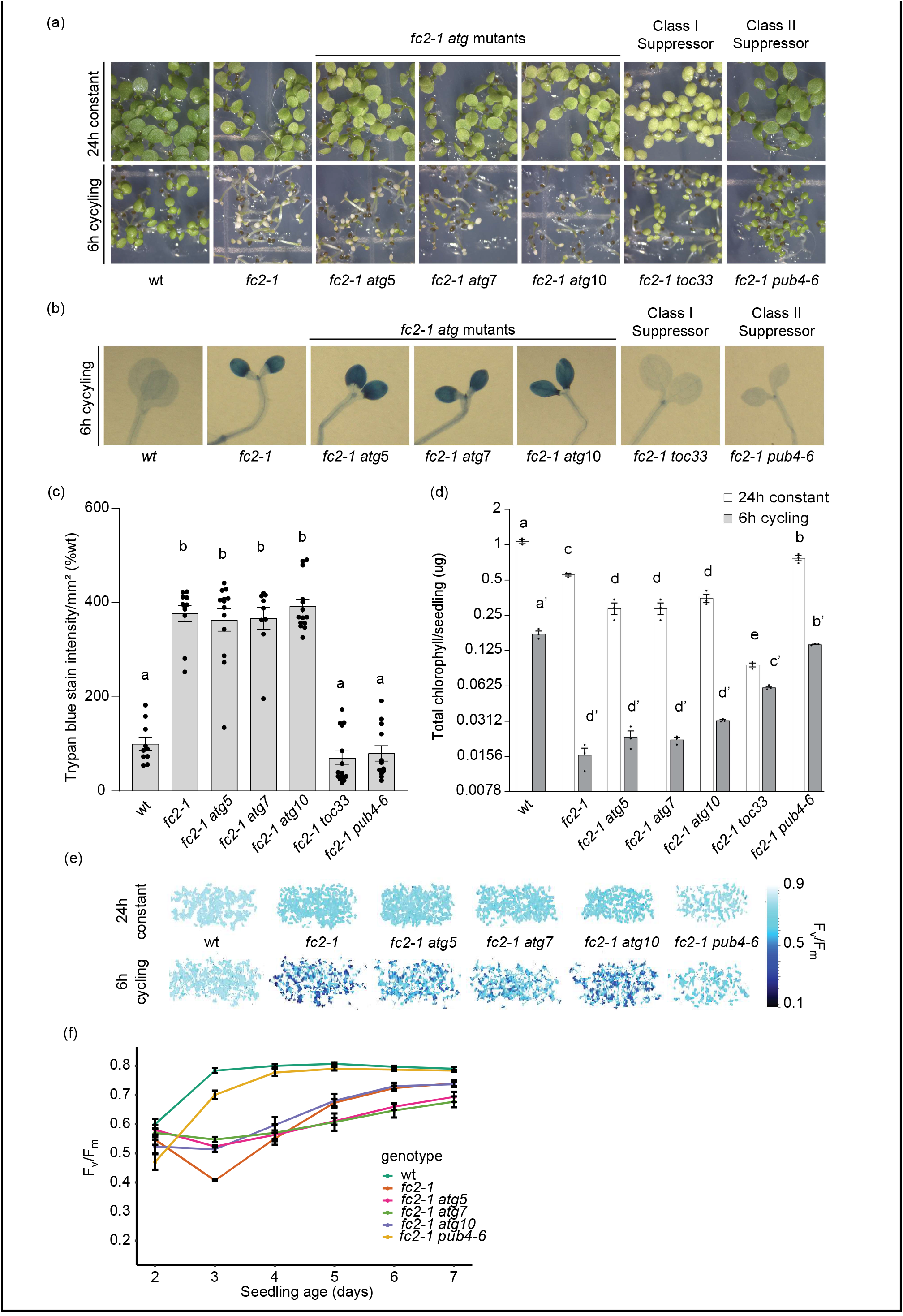
Blocking macroautophagy does not suppress ^1^O_2_-induced cell death in *fc2* seedlings. *atg* mutations were tested for their ability to suppress cell death in the *fc2-1* mutant. **A)** Seven-day old seedlings grown in constant light (24h) or 6h light/18h dark (6h) cycling light conditions. **B)** Trypan blue stains of seedlings from panel A. The dark blue color is indicative of cell death. **C)** Mean values (+/- SEM) of the trypan blue signal in panel B (n ≥ 10 seedlings). **D)** Mean chlorophyll content (+/- SEM) of six-day old seedlings grown in 24h light or 6h cycling light conditions (n = 3 biological replicates). **E)** Representative images of maximum quantum yield of PSII (F_v_/F_m_) measured from three-day-old seedlings grown in the indicated light regiment. **F)** Changes in F_v_/F_m_ of seedlings grown over seven days in 6h cycling light conditions. Mean values are shown +/- SEM (n = 3 biological replicates). Statistical analyses were performed by one-way AVOVA tests followed by Tukey’s HSD. Different letters above bars indicate significant differences (p value ≤ 0.05). For panel D, separate statistical analyses were performed for the different light treatments and the significance for the 6h cycling values is denoted by letters with a ’. In all bar graphs, closed circles represent individual data points.

Additionally, the low chlorophyll content of *fc2-1* seedlings grown under cycling light is not significantly affected by the three *atg* mutations, further demonstrating their inability to suppress the *fc2-1* cell death phenotype (Fig. 3d). Compared to *fc2-1*, however, *fc2-1 atg* double mutants have a significantly lower level of chlorophyll in 24h constant light conditions, which may indicate a role for autophagy in chloroplast development or stress responses. This effect on chlorophyll content is partially independent in the *fc2-1* background, as single *atg* mutants have reduced chlorophyll content compared to wt (Fig. S2d). Together, these data demonstrate *atg* mutations do not suppress the *fc2-1* cell death phenotype, indicating core autophagy proteins ATG5, ATG7, and ATG10 are dispensable in ^1^O_2_-mediated cellular degradation at the seedling stage.

To further investigate the effect of *atg* mutations on the physiology of *fc2-1*, we next measured maximum quantum yield of PSII (F_v_/F_m_), an indicator of photoinhibition and plant stress (50), in the same seedlings. Under 24h constant light conditions, photosynthetic efficiency is slightly delayed in *fc2-1* mutants compared to wt, but gradually increases to wt levels after five days (Fig. 3e and S3a). Under 6h light/18h dark cycling light conditions, however, photosynthetic efficiency in *fc2-1* mutants decreases at day three and then gradually recovers over the next four days (Fig. 3e and f). This likely reflects the level of cellular degradation occurring in these mutants.

While the cell death suppressor *pub4-6* mostly restored photosynthetic efficiency to the *fc2-1* mutant under cycling light conditions, the three *atg* mutations did not (Fig. 3e). However, a closer look at the time course analysis shows that the three *fc2-1 atg* double are slightly less affected by ^1^O_2_ stress at day three compared to *fc2-1* (Fig. 3f), but then recover more slowly than *fc2-1* over the following four days. This effect is specific to cycling light as *fc2-1 atg* double mutants are nearly identical to *fc2-1* in 24h constant light (Fig. S3a). Moreover, this effect is specific to the *fc2-1* background, as single *atg* mutants are almost indistinguishable from wt in either condition (Fig. S3b-d). Together, these results further support macroautophagy does not play a positive role in ^1^O_2_–induced chloroplast or cell degradation in the *fc2-1* mutant. However, the complex effect of the *atg* mutations on photosynthetic efficiency during ^1^O_2_ stress, suggests macroautophagy may play a role in maintaining photosynthetic efficiency under photo-oxidative stress.

### Blocking macroautophagy does not suppress cell death or growth inhibition in *fc2* adult plants

Chlorophagy has previously been shown to occur in adult plants after UVB and EL stress (34,35). Therefore, we tested if macroautophagy may play a stage-specific role in plant development by assessing the phenotypes of *fc2-1 atg* double mutant adults plants. To do this, we grew plants for two weeks in 24h constant light and then transferred them to 16h light/8h dark cycling light conditions. As a control, another set of plants remained under 24h constant light for a third week. While *fc2-1* plants are healthy even after three weeks in 24h constant light, they exhibit necrotic lesions in leaves and reduced growth (dry weight biomass) after one week in cycling light (Fig. 4a and b). These necrotic lesions were confirmed to be areas of cell death by trypan blue staining of leaves (Fig. 4c and d). When mature two-week-old *fc2-1 atg* double mutants were exposed to cycling light stress, all three had a similar amount of leaf necrosis, cell death, and impaired growth compared to *fc2-1* single mutants. The three single *atg* mutants behaved similarly to wt indicating the stress phenotypes described above are dependent on the *fc2-1* background (Fig. S4a-d). Together, these data suggest ^1^O_2_-induced chloroplast and cell degradation is not dependent on macroautophagy in either the seedling or adult stages.

**Figure 4.**
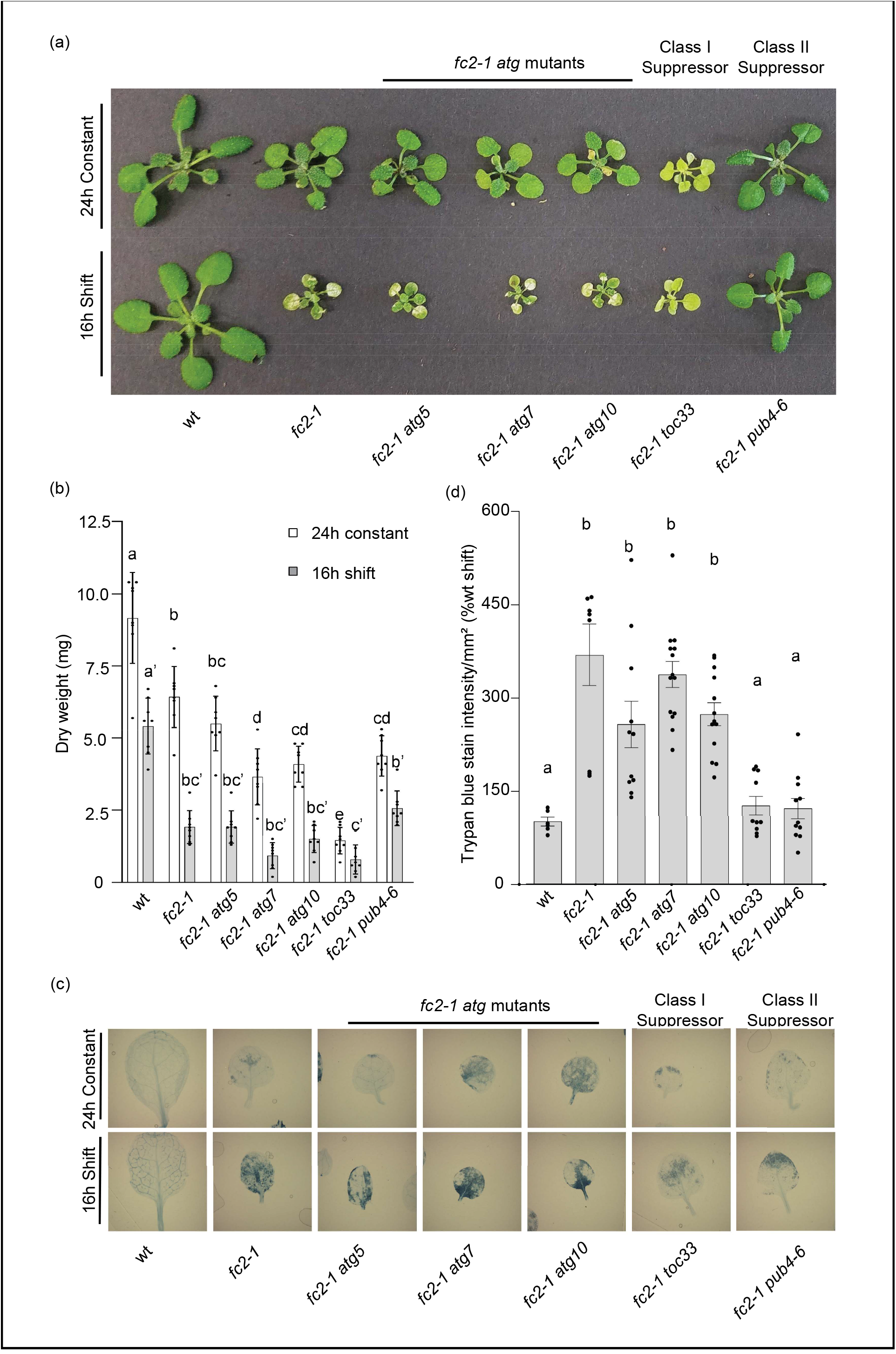
Blocking macroautophagy does not suppress ^1^O_2_-induced cell death in *fc2* adult plants. The phenotypes of *fc2-1 atg* double mutant plants were assessed in the adult stage. **A)** Three-week old plants grown in 24h constant light or under stressed conditions (two weeks in 24h constant light and one week in 16h light/8h dark cycling light conditions). **B)** Mean biomass (+/- SD) of same plants (n = 8 plants). **C)** Representative trypan blue stains of single leaves from same plants. Dark blue color is indicative of cell death. **D)** Quantification of mean trypan blue signal (+/- SEM) in panel C (n ≥ 6 leaves from individual plants). Statistical analyses were performed by one-way AVOVA tests followed by Tukey’s HSD. Different letters above bars indicate significant differences (p value ≤ 0.05). For panel B, separate statistical analyses were performed for the different light treatments and the significance for the light stressed group is denoted by letters with a ’. In all bar graphs, closed circles represent individual data points.

### Macroautophagy is not required for transmitting the ^1^O_2_ retrograde signal in *fc2* mutants

In addition to chloroplast degradation and cell death, ^1^O_2_ also leads to retrograde signals that regulate hundreds of genes in the nucleus (9). Under cycling light, stressed *fc2* mutants have been shown to have a clear induction of multiple nuclear stress marker genes (specific ^1^O_2_ response genes; *BAP1* (*AT3G61190*) and *ATPase* (*AT3G28580*) (8), general ROS response genes; ZAT12 (*AT5G59820*) and *CYC81D8* (*At4g37370*) (51), and genes specific to stressed *fc2-1* seedlings; SIB1 (*AT3G56710*) and *HSP26*.*5* (*AT1G52560*) (14)).

As there is no apparent suppression of the physiological phenotypes of stressed *fc2-1* in the *fc2-1 atg* double mutants, we next asked if core autophagy mutations lead to changes in the induction of these key stress markers. To do this, we extracted RNA from four-day-old seedlings grown under 6h light/18h dark cycling light conditions and used RT-qPCR to monitor transcript levels of these six marker genes. As expected, the *fc2-1* mutant significantly induced all six genes compared to wt (Fig. 5). The three *fc2-1 atg* double mutants also induced all six genes compared to wt and were virtually indistinguishable from the single *fc2-1* mutant. Together, these data suggest that core autophagy proteins are not necessary for the chloroplast retrograde signals induced in *fc2-1* seedlings.

**Figure 5.**
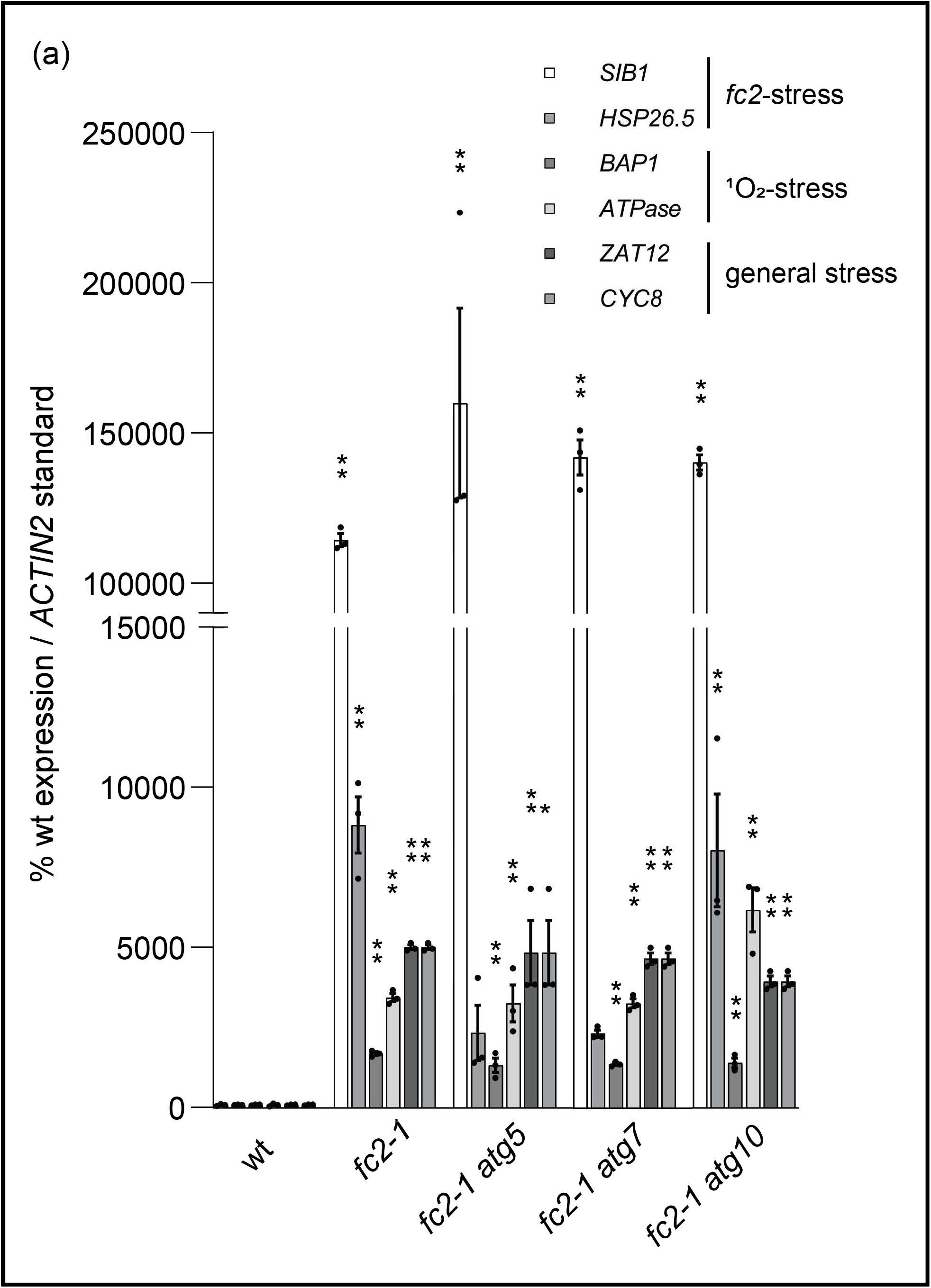
Macroautophagy is not required for ^1^O_2_ retrograde signaling in *fc2* mutants. The expression of nuclear genes controlled by chloroplast ^1^O_2_ stress signaling were monitored in the *fc2-1 atg* double mutants. **A)** RT-qPCR analyses of stress gene transcript abundance in four-day old *fc2-1* and *fc2-1 atg* double mutants grown under 6h light/18h dark cycling light conditions collected one hour after subjective dawn. Shown are means of biological replicates (n = 3) +/- SEM. Statistical analyses were performed by a one-way ANOVA followed by Dunnett’s multiple comparisons test with the wt. * and ** indicates an adjusted p value of ≤ 0.05 and ≤ 0.01, respectively. Closed circles represent individual data points.

### Chloroplast degradation in the *fc2-1* mutant is not dependent on macroautophagy

The above results suggest the *atg* mutations do not lead to *fts* phenotypes and cannot block cellular degradation in the *fc2-1* mutant background under light cycling conditions. To test if the ATG proteins and macroautophagy play a specific role is dismantling ^1^O_2_-damaged chloroplasts in *fc2-1*, we used transmission electron microscopy to image four day old seedlings grown under permissive 24h constant light conditions. Under such conditions, *fc2-1* mutants avoid cell death, but a subset of chloroplasts are still degraded and can sometimes be observed protruding or blebbing into the central vacuole (14).

As expected, most of the chloroplasts in *fc2-1* seedlings appeared intact, but a subset were in the process of being degraded (Fig. 6a and b) (18.6% of the total chloroplasts imaged). These degrading chloroplasts can be identified by their abnormal shapes, severely disorganized internal membranes, and compressed thylakoids. Most chloroplasts in *fc2-1 atg5* and *fc2-1 atg7* mutants also appeared intact, but a larger percentage were being degraded (39.0% and 43.5% in *fc2-1 atg5* and *fc2-1 atg7*, respectively) (Fig. 6a and b). In some cases, chloroplasts in these double mutants could be observed protruding or blebbing into the central vacuole (Fig. 6c and d) similar to what has previously been observed in single *fc2-1* mutants (14). Such structures were observed two times in 15 *fc2-1 atg5* cells, two times in 10 *fc2-1 atg7* cells, and zero times in 15 *fc2-1* cells. Together these results suggest *atg5* and *atg7* are unable to block selective chloroplast degradation or chloroplast blebbing in the *fc2-1* mutant. No chloroplast degradation was observed in wt, *atg5*, or *atg7*.

**Figure 6.**
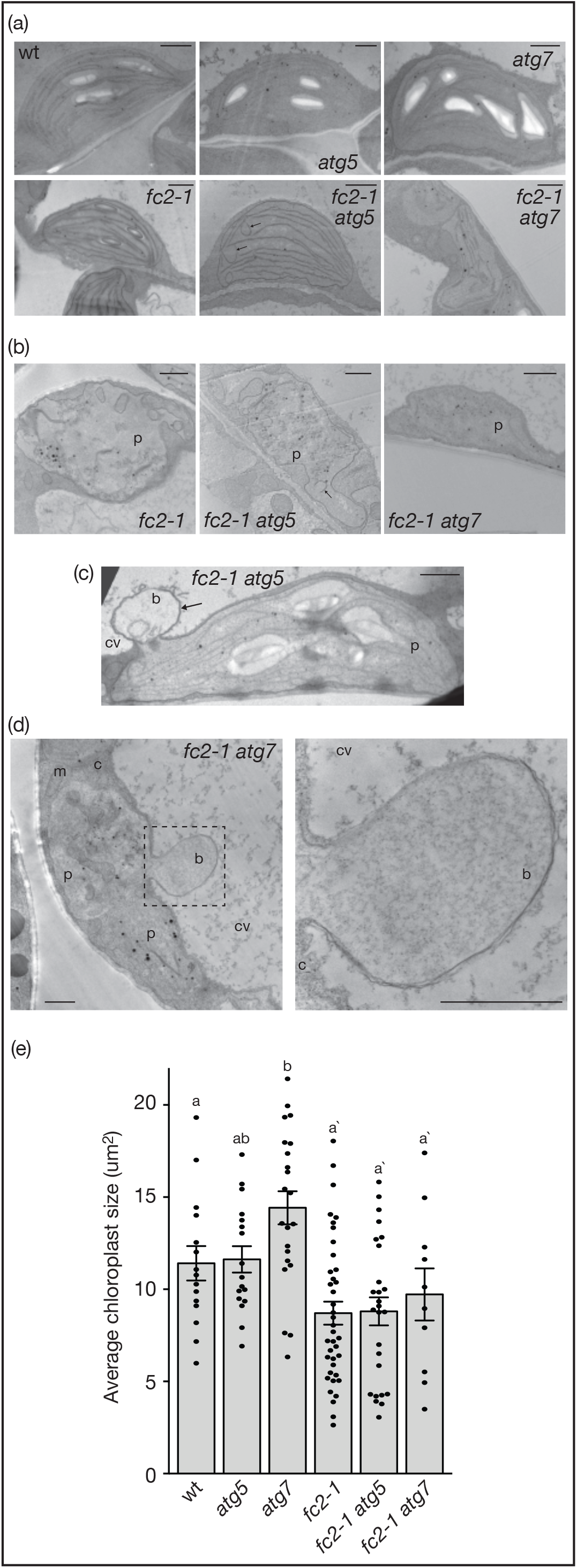
Macroautophagy is not required for selective chloroplast degradation in in *fc2* mutants. Chloroplast ultrastructure was assessed by transmission electron microscopy (TEM) in the *atg* mutants. Shown are representative **A)** intact and **B)** degrading chloroplasts in four day old seedlings grown under 24h constant light conditions. Arrows indicate unusual horseshoe membrane structures. Shown is a blebbing chloroplast interacting with the central vacuole in the **C)** *fc2-1 atg5* and **D)** *fc2-1 atg7* mutants. Arrows indicate the bleb-like structure protruding into the central vacuole. The right panel is a zoomed-in image of the boxed region. b; bleb, c; cytoplasm, cv; central vacuole, m; mitochondria, p; plastid. Bars = 1 µm. **E)** Mean chloroplast area (+/- SEM) from the same seedlings (n ≥ 10). Statistical analyses were performed using one-way AVOVA tests followed by Tukey’s HSD. Different letters above bars indicate significant differences (p value ≤ 0.05). Separate statistical analyses were performed for seedlings in the wt and *fc2-1* backgrounds and the significance for the *fc2-1* background group is denoted by letters with a ’. Closed circles represent individual data points.

In addition to increasing the percentage of degrading chloroplasts, the *atg* mutations also affected chloroplast ultrastructure in the *fc2-1* background. In many cases, *fc2-1 atg5* and *fc2-1 atg7* chloroplasts had irregular shapes or contained unusual horseshoe shaped membranes/vesicles (Fig. 6a and b). Such abnormalities were not observed in the single *atg5* and *atg7* mutants. The average chloroplast area was not reduced by the *atg* mutations in either the wt or *fc2-1* backgrounds, suggesting chloroplast development was not grossly delayed by the loss of macroautophagy (Fig. 6e). As such, macroautophagy may still play a role in maintaining chloroplast structure and/or function during photo-oxidative stress.

### Genes encoding microautophagy-related proteins are induced in stressed *fc2* mutants

The observation that ^1^O_2_-mediated chloroplast degradation and blebbing into the central vacuole still occurs in *fc2-1 atg* double mutants suggests canonical autophagosome formation is dispensable for such a degradation pathway. This led us to consider alternative forms of autophagy that may play a role in ^1^O_2_-mediated selective chloroplast degradation. As the chloroplast blebbing observed in this process appears to involve invagination into the central vacuole, we aimed to investigate if microautophagy may play a role in this mechanism. Indeed such structures were also observed in *fc2-1 atg* double mutants that appeared independent of chloroplasts (Fig. 7a), suggesting microautophagy is still active in these lines. However, quantification of microautophagy or chloroplast blebbing events is difficult due to their infrequency in two-dimensional micrographs (14). This is likely due to the small size of the connection point between the mircoautophagosome structure and the cytoplasm.

**Figure 7.**
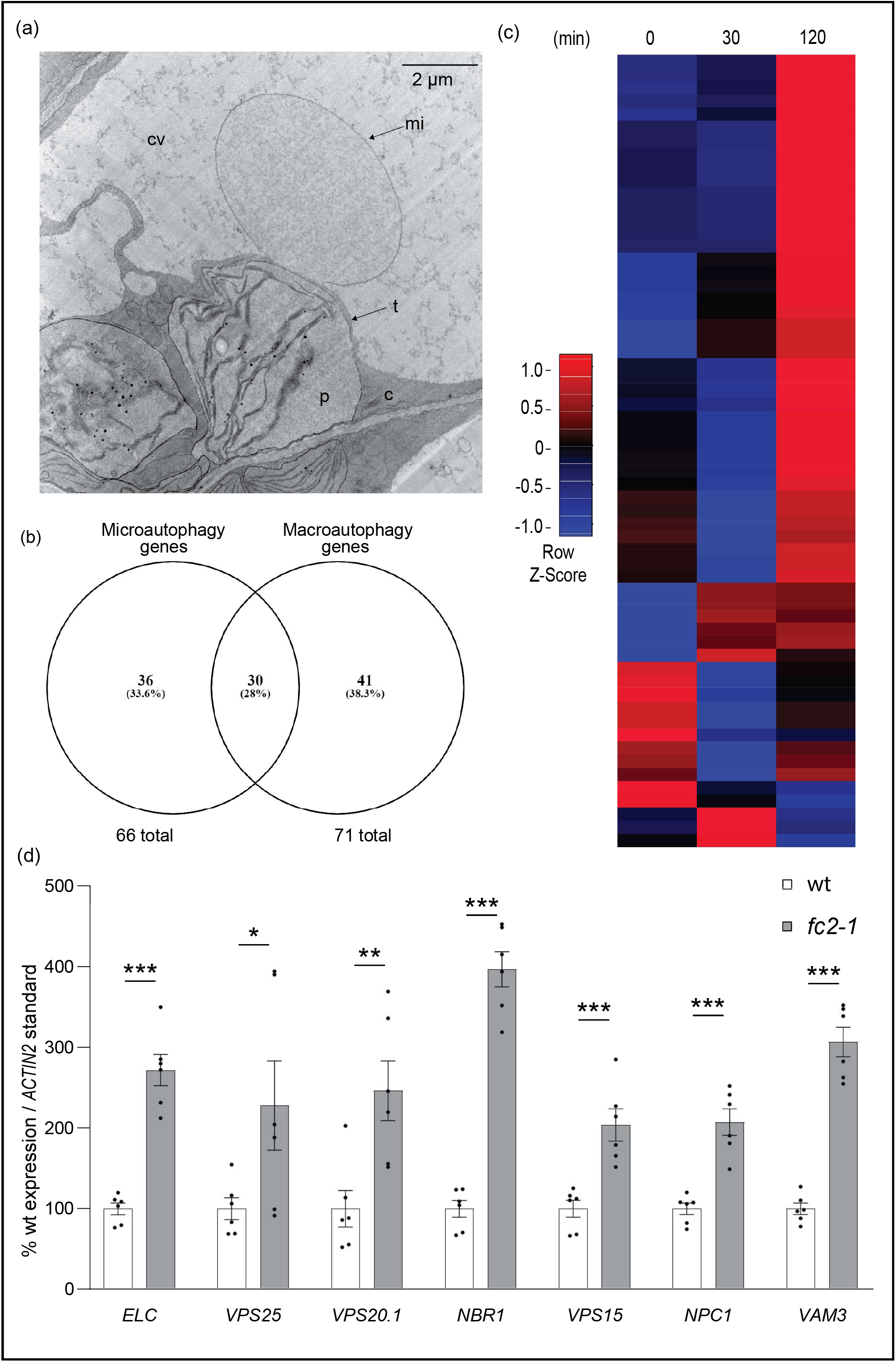
Microautophagy-related genes are transcriptionally induced in stressed *fc2* seedlings. The induction of microautophagy was assessed in *fc2-1* seedlings. **A)** Shown is a representative TEM image of the invagination of the vacuolar membrane associated next to a degrading chloroplast in the *fc2-1 atg5* mutant. c; cytoplasm, cv; central vacuole, mi; microautophagy structure, p; plastid, t; tonoplast. **B)** A Venn diagram showing overlap of putative microautophagy-related genes and autophagy-related genes used in this study (Tables S1 and S2). **C)** A heatmap of microautophagy-related gene expression (relative to wt) in etiolated seedlings at time points prior to (0 min) and during de-etiolation (30 min and 120 min). Microarray data was previously generated from etiolated seedlings grown in the dark for four days at time points before and after light exposure (14). The list of genes considered is presented in Table S2. **D)** RT-qPCR analysis of select microautophagy-related transcripts from four-day-old seedlings grown under 6h light/18h dark light cycling conditions. Shown are mean values +/- SEM (n = 6 biological replicates). Statistical analyses were performed by student’s t-tests. *, **, *** indicate a p-value of ≤ 0.05, ≤ 0.01, and ≤ 0.001, respectively. In all bar graphs, closed circles represent individual data points.

Therefore, to determine if microautophagy may be responsible for structures associated with chloroplast degradation in *fc2-1* mutants, we monitored the expression of genes that have been predicted to play a role in microautophagy due to their homology to microautophagy and microautophagy-related processes in *Saccharomyces cerevisiae* (31). From this work, we generated a list of 66 genes to investigate in *fc2-1* (Table S2). As microautophagy itself is not well defined, particularly in plants (31), we first determined which genes this microautophagy gene list were shared with our manually curated list of general autophagy genes used in Fig. 1a and Table S1. There were 30 genes common to both lists, nearly all of which were the core ATG genes (Fig. 7b). As most of the predicted microautophagy-related genes were unique, we generated a heatmap using the same previously published microarray data set of wild type (wt) and *fc2-1* seedlings before and during de-etiolation (14). Again, we saw a pattern of induction in this subset of predicted microautophagy-related genes, primarily 120 minutes after de-etiolation (79% of genes, with no cutoffs applied) (Fig. 7c). As these microarray data were from seedlings prior to and during de-etiolation, we again chose to validate these patterns in stressed four day-old *fc2-1* seedlings using RT-qPCR. Here, we probed select genes from a number of functional gene categories (31): ESCRT-I, *AT3G12400* (*ELC*); ESCRT-II, *AT4G19003* (*VPS25*); ESCRT-III, *AT5G63880* (*VPS20*.*1*); Vacuolar Selective Mono-Ub Receptor, *AT4G24690* (*NBR1*); PI3K Complex, *AT4G29380* (*VPS15*); Niemann–Pick type C proteins (NPC), *AT1G07230* (*NPC1*); SNARE, *AT5G46860* (*VAM3*). In all cases, transcript abundance was significantly increased in *fc2-1* compared to wt. Together, these data suggest microautophagy is transcriptionally induced in *fc2-1* seedlings under ^1^O_2_-generating conditions. Based on the type of structures known to associate with ^1^O_2_-stressed chloroplasts, this may indicate a putative role for microautophagy in the mechanism by which ^1^O_2_-damaged chloroplasts are degraded in *fc2* mutants.

## Discussion

Chloroplast ROS accumulates under various environmental stresses and has the ability to initiate multiple signaling pathways. Usually, however, more than one type of ROS is generated simultaneously and it is difficult to understand the signaling contributions of each species (1,52). As such, the use of mutants that specifically and conditionally accumulate only one type of ROS have been very important for understanding the signaling abilities and mechanisms of individual ROS. The *fc2* mutant, that specifically accumulates chloroplast ^1^O_2_ under diurnal signaling conditions, has been an extremely useful system to dissect the role of this ROS in initiating chloroplast quality control and cellular degradation pathways. Genetic suppressor screens have identified chloroplast gene expression and the cellular ubiquitination machineries as playing important roles in these processes, providing insight into how ^1^O_2_ initiates a stress signal and how individual chloroplasts may be recognized for turnover (14,15). While such chloroplast degradation appears to be a targeted and orderly process, the cellular machinery involved has remained obscure. In order to further understand the mechanisms behind this degradation, here we tested the possibility that autophagy is involved in ^1^O_2_–mediated cell death pathways or in removing ^1^O_2_–damaged chloroplasts.

Previous work has shown autophagy can be involved in translocating UVB or EL-damaged chloroplasts to the central vacuole in a process referred to as chlorophagy (33,35). The autophagosome/chloroplast interaction differed under the two types of stresses. UVB caused a classic macroautophagy-like response with full envelopment of the damaged chloroplasts by the autophagosome (33). EL, on the other hand, involved a microautophagy-like response with the autophagosome partly surrounding the damaged chloroplast, which was engulfed by the tonoplast (35). In both cases, chlorophagy were entirely dependent on the core autophagy machinery and chloroplasts remained in the cytoplasm in *atg5* and *atg7* mutants. However, when the same mutations (including a third mutation affecting core autophagy protein ATG10) were introduced into the *fc2* background, they did not suppress ^1^O_2_–induced cellular degradation. All three *fc2 atg* double mutants still experienced cell death as seedlings or adult plants when grown under diurnal cycling light (Figs. 3a and 4a). Furthermore, the chloroplast protrusion or blebbing into the central vacuole, a characteristic of ^1^O_2_-mediated selective chloroplast degradation in *fc2* (14), was still present in *fc2 atg5* and *fc2 atg7* double mutants when grown under permissive 24h constant light. Taken together, these results demonstrate that chlorophagy is not directly involved in ^1^O_2_-induced chloroplast quality control pathways or cell death. Instead, the core autophagy machinery and macroautophagy was dispensable.

While selective chloroplast degradation can occur in *fc2* mutants without core autophagy machinery, there still appears to be a relationship between the two. Under diurnal cycling light conditions that produce chloroplast ^1^O_2_, *fc2* seedlings respond by transcriptionally inducing autophagy-related genes, including those encoding core autophagy proteins (Fig. 1a and d). Autophagosomes containing ATG8a are also present in these cells, even though they do not appear to be directly interacting with chloroplasts for degradation (Fig. 2). Instead, such a phenotype may be related to an induced starvation response in *fc2* seedlings as evidenced by the induction of carbon and nitrogen starvation marker genes (Fig. 1e). It is possible the chloroplast dysfunction and cellular degradation in ^1^O_2_-stressed *fc2* seedlings has a pleotropic effect on cellular metabolism leading to the induction of autophagosomes and the remobilization of nutrients. As such, autophagomes and the induction of general autophagy genes may be partly a consequence rather than the cause of the cellular degradation observed in *fc2* seedlings.

While the *atg* mutations did not suppress cellular degradation, they did affect how *fc2* chloroplasts responded to photo-oxidative stress. When grown for three days under light cycling conditions, the *atg5, atg7*, and *atg10* mutations appeared to slightly protect the *fc2* mutant from photoinhibition (F_v_/F_m_). However, compared to *fc2-1*, photosynthesis in the double mutants was also slower to recover over time (Fig. 3e, and 3f). These effects on photosynthesis were dependent on the *fc2-1* background, as *atg* single mutants were almost indistinguishable from wt. Such a result suggests the loss of autophagy may have altered the sensitivities of chloroplasts to photo-oxidative damage. Indeed, an analysis of chloroplast ultrastructures demonstrated that chloroplasts in the double mutants were irregular in shape and contained unusual internal membrane structures (Fig. 6a and b). The rate of chloroplast degradation also appeared to increase in the double mutants compared to *fc2-1*. As such, autophagy may still be important for maintaining chloroplast homeostasis and function during ^1^O_2_ stress. This could be due for indirect reasons (e.g., nutrient remobilization or general cell health) or perhaps by directly regulating chloroplast proteins levels through autophagy-dependent protein quality control pathways. Such pathways can involve RuBisCO-containing bodies (RCB’s) (53) and ATG8-interacting protein (ATI) bodies (54) that package chloroplast proteins for degradation in the central vacuole in a piecemeal fashion.

If chlorophagy and macroautophagy are not directly involved in ^1^O_2_–induced chloroplast degradation, then what process is being used? Compared to chlorophagy, chloroplast degradation in *fc2* cells is visually distinct. Chloroplasts in *fc2* mutants already appear to be in a state of degradation before they protrude or “bleb” into the central vacuole without the obvious presence of autophagosome membranes (Fig. 6d and (14)). Such a structure is visually similar to microautophagy, involving the invagination of the central vacuole around cytosolic components (31,55). Unlike EL-induced chlorophagy, this type of microautophagy is independent of autophagosomes and the core autophagy machinery. An expression analysis of predicted microautophagy-related genes showed a clear pattern of induction in the *fc2* mutant (Fig. 7d). As such, microautophagy is a promising candidate mechanism for selective vacuolar transport of ^1^O_2_-damaged chloroplasts in *fc2* mutants, a possibility to be further explored.

## Conclusions

In this work, we have demonstrated that ^1^O_2_-induced cellular degradation and selective chloroplast degradation in the *fc2* mutant occurs via a process distinct from chlorophagy and does not require macroautophagy or the core autophagy machinery. This is supported by a previous report demonstrating the PUB4 protein, which is required for selective chloroplast degradation in the *fc2* mutant (14), is dispensable for chlorophagy after EL stress (36). Together, these results suggest alternate pathways of chloroplast degradation can be used in the cell, possibly depending on the type of stress or damage being experienced. Such a model would explain the altered starvation responses and severe early senescence phenotypes of *pub4 atg5* and *pub4 atg7* double mutants that may lack two independent pathways for dismantling chloroplasts (36). Further work will still be necessary to understand the different roles of these pathways during stress, how they recognize damaged chloroplasts, and how these chloroplasts are ultimately transported to the central vacuole. A deeper understanding of the mechanisms behind these pathways will be of central importance in understanding how cells maintain chloroplast function and homeostasis under dynamic and stressful conditions and may help to provide molecular tools to control photosynthesis and yield in agriculturally important crop species.

## Methods

### Biological material, growth conditions, and treatments

*Arabidopsis thaliana* ecotype *Columbia* (Col-0) was the wild type (wt) line and was used to generate all transgenic constructs. T-DNA lines GABI_766H08 (*fc2-1*) and GABI_655B06 (*atg7-2*) from the GABI-Kat collection (56), SALK_084434 (*atg10-1*) from the SALK (The Salk Institute Genomic Analysis Laboratory) collection (57) and SAIL_129_B07 (*atg5-1*) from the SAIL collection (58) were described previously (30,46,48,59). The point mutants *pub4-6* and *toc33* were described previously (14). Additional information is listed in Table S3. Double mutant lines were generated via crossing and confirmed by extracting genomic DNA (following a CTAB-based protocol (60)) and confirming genotypes using PCR-based markers (primers listed in Table S4).

Seeds used in growth experiments were sterilized overnight using chloride gas and then resuspended in a 0.1% agar solution. Resuspended seeds were then spread on plates containing Linsmaier and Skoog medium pH 5.7 (Caisson Laboratories North Logan, UT) in 0.6% micropropagation type-1 agar powder. Plates were then stratified for three-five days and then germinated and grown in constant light (24h day) or diurnal cycling light (16h day/8h night) conditions at a light intensity of ∼100 photons µmol m^-2^ sec^-1^ at a temperature of 21°C. For adult plant experiments, seven-day old seedlings grown in constant light conditions were carefully transferred to soil and growth was continued under similar conditions in a reach-in plant growth chamber. For dark starvation experiments, plates containing four day old seedlings were wrapped in aluminum foil three times and then placed back into the growth chamber. Six days later, these plates were opened under dim light and the seedlings were imaged by confocal microscopy. Photosynthetically active radiation was measured using a LI-250A light meter with a LI-190R-BNC-2 Quantum Sensor (LiCOR).

*Agrobacterium tumefaciens* was grown in liquid Miller nutrient broth or solid medium containing 1.5% agar (w/v). Cells were grown at 28°C with the appropriate antibiotics and liquid medium was shaken at 225 rpm.

### Construction of transgenic *Arabidopsis* lines

The vector containing the *UBQ10*::*GFP-ATG8a* construct (At4g21980.1 (61)) was transformed into the *A. tumefaciens* strain GV301, which was subsequently used to transform *Arabidopsis* via the floral dip method. T_1_ plants were selected for their ability to grow on Basta-soaked soil and propagated to the next generation. T_2_ lines were monitored for single insertions (segregating 3:1 for Basta resistance:sensitivity) and propagated to the next generation. Finally, homozygous lines were selected in the T_3_ generation based on 100% Basta-resistance

### Biomass measurements

For dry weight (DW) biomass measurements, eight replicates of each line and growth condition were weighed then stored in envelopes for desiccation. Tissue was desiccated in a 65°C oven for 48 hours and then remeasured to record dry weight.

### Chlorophyll fluorescence measurements

To measure Maximum PSII quantum yield (F_v_/F_m_), seedlings were germinated under 24h constant or 6h/18h light/dark cycling light conditions. F_v_/F_m_ was measured two hours after subjective dawn every day for six days. Plates were placed in the FluorCam chamber (Closed FluorCam FC 800-C/1010-S, Photon Systems Instruments) for 15 minutes to dark acclimate without a plate cover, and then the measurements were taken following the manual provided by the manufacturer. Each genotype was sown in triplicate, on separate plates, and F_v_/F_m_ measurements were taken from each replicate consecutively.

### Chlorophyll measurements

Total chlorophyll was extracted using 100% ethanol from seven-day-old seedlings. Cell debris was pelleted thrice at 12,000 x g for 30 min at 4 °C. Chlorophyll was measured spectrophotometrically in 100 μl volumes in a Biotek Synergy H1 Hybrid Reader and path corrections were calculated according to (62). Chlorophyll levels were normalized per seedling (∼ 50-130 per line and counted prior to germination).

### Cell death measurements

Cell death was assessed using trypan blue staining as previously described (15). Briefly, seedings were collected at seven days and then transferred to a solution of Trypan Blue (glycerol: 1.25 ml, trypan blue solution: 0.625 ml, ddH_2_O: 0.625 ml, lactic acid: 1.25 ml, phenol: 1.25 ml, 100% ethanol: 10 ml). The solution was then boiled for 1 minute, incubated overnight, and then destined with a solution of saturated chloral hydrate (2.5g per 1 ml water) Chloral hydrate was then removed, and seedlings were stored in 30% (v:v) glycerol solution prior to imaging on a dissection scope. For each replicate, staining intensity was measured across one entire cotyledon (seedlings) or leaf (adult plants) using ImageJ software.

### RNA extraction and RT-qPCR

Total RNA was extracted from whole seedlings using the RNeasy Plant Mini Kit (Qiagen) and cDNA was synthesized using the Maxima first-strand cDNA synthesis kit for RT-qPCR with DNase (Thermo Scientific) following the manufacturer’s instructions. Real-time PCR was performed using the SYBR Green Master Mix (BioRad) with the SYBR Green fluorophore and a CFX Connect Real-Time PCR Detection System (Biorad). The following 2-step thermal profile was used in all RT-qPCR: 95 °C for 3 min, 40 cycles of 95 °C for 10s, and 60 °C for 30s. All expression data was normalized according to *ACTIN2* expression. All primers used for RT-qPCR are listed in Table S4.

### Transmission electron microscopy

Four day old seedlings were prepared and imaged by transmission electron microscopy as previously described (15). Chloroplast area was analyzed using ImageJ.

### In vivo protein localization

Four day-old seedlings (or 10 day old seedlings for dark starvation experiments) were imaged on a Zeiss 880 inverted confocal microscope. The excitation/emissions wavelengths used were 458/633 for chlorophyll autofluorescence and 488/510 for GFP. Resulting images were processed using Zen Blue (Carl Zeiss) software and analyzed in Adobe PhotoShop and ImageJ/Fiji.

### Gene expression meta-analysis

To monitor the expression of autophagy-related genes controlled by chloroplast ^1^O_2_ signals, we used a previously published data set of a microarray analysis of four-day-old etiolated wt and *fc2-1* seedlings before and after (30 and 120 minutes) light exposure (120 µmol photons m^-2^ sec^-1^) (14). Manually curated list core autophagy and autophagy related genes (Table S1) were compiled from gene ontology biological process term GO:0016238 from the Arabidopsis Information Resource (TAIR) website (https://www.arabidopsis.org/). Parent (24 loci) and children terms (60 loci) were collected. Duplicate and undefined loci were removed. A list of putative microautophagy-related genes (Table S2) and associated loci were compiled from (31), and additional information was obtained from TAIR. A data matrix reflecting the microarray expression data was then compiled for each of these lists. Genes with no detected expression data or no associated microarray probe were removed from the data matrix. Heatmaps were generated using the heatmapper package (https://github.com/WishartLab/heatmapper) on http://www.heatmapper.ca/ (63). Average linkage was used for the clustering method and Pearson correlation was used for the distance measurement method. Venn diagrams were generated using Venny Diagram (https://bioinfogp.cnb.csic.es/tools/venny/) (64).

## List of abbreviations

^1^O_2_: Singlet oxygen,
QC: Quality control,
ROS: Reactive oxygen species,

## Declarations

### Ethics approval and consent to participate

Not applicable

### Consent for publication

Not applicable

### Availability of data and materials

All data generated or analyzed during this study are included in this published article (and its supplementary information files). One exception is the previously published microarray data set from Woodson et al. 2015 that is deposited at Gene Expression Omnibus (GSE71764).

### Authors’ contributions

MDL, KEF, MAK, DWT, and JDW planned and designed the research. MDL performed all RT-qPCR, physiological growth experiments, and plant treatments. KEF performed all microscopy experiments and their sample preparations. MAK developed and performed all chlorophyll fluorescence experiments. DWT developed and performed chlorophyll assays. JDW conceived the original scope of the project and managed the project. MDL and JDW wrote the manuscript. All authors contributed to data analysis, collection, and interpretation. All authors reviewed the manuscript and approved the final version.

## Acknowledgments

The authors thank Dr. Masanori Izumi (Riken) for sharing the *UBQ10*::*GFP-ATG8a* construct, Dr. David Baltrus (U. Arizona) for providing a plate reader for chlorophyll measurements, and Kamran Alamdari (U. Arizona) for technical assistance in generating transgenic *Arabidopsis* plants.

## Short legends for supporting information

### Supplemental Figures

Figure S1. Validation of *atg5, atg7*, and *atg10* null mutations in the *fc2-1* background.

Figure S2. Phenotypes of *atg5, atg7*, and *atg10* single mutant seedlings.

Figure S3. Assessment of photosynthetic efficiency in the *atg* single and *fc2 atg* double mutants.

Figure S4. Phenotypes of *atg5, atg7*, and *atg10* single mutant adult plants.

### Supplemental Tables

Table S1. List of autophagy-related genes and expression data used in heatmap generation.

Table S2. List of predicted microautophagy-related genes and expression data used in heatmap generation.

Table S3. Mutants used in this study Table S4. Primers use in this study

