## Supplemental figures and tables for "Singlet oxygen-dependent chloroplast degradation is independent of macroautophagy in the *Arabidopsis ferrochelatase* two mutant"

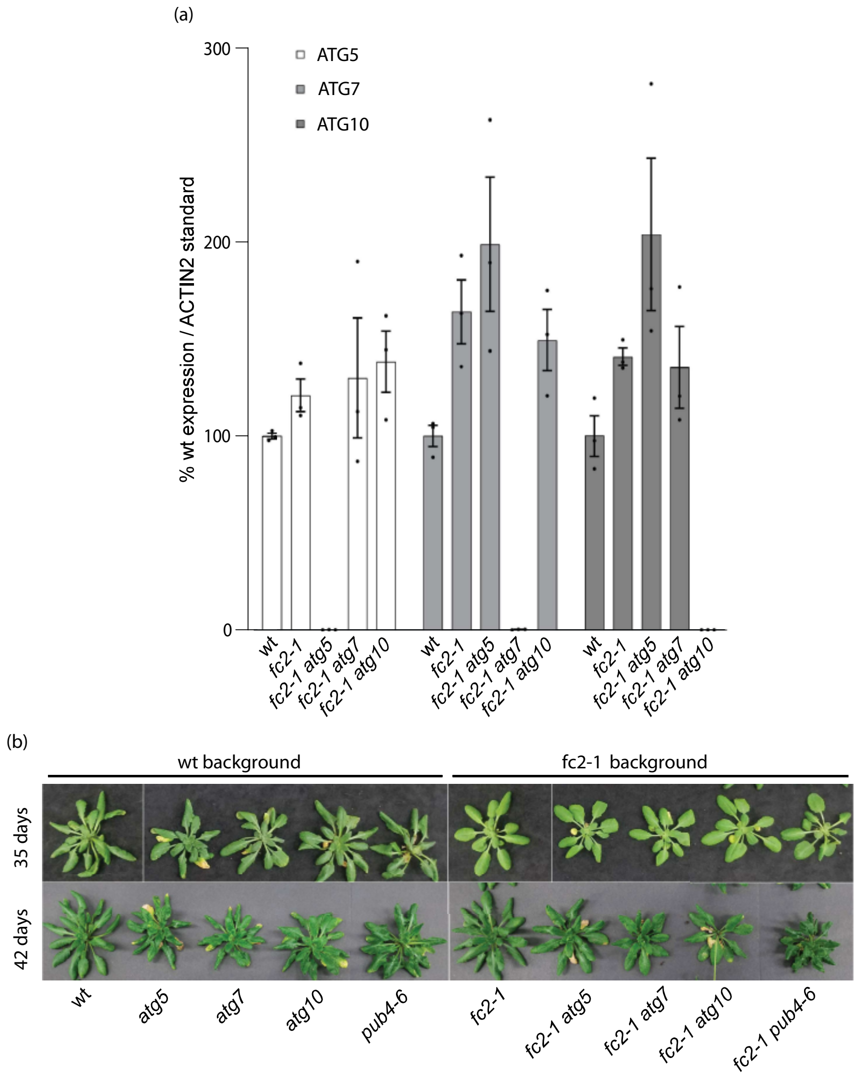


Figure S1. Validation of *atg5*, *atg7*, and *atg10* null mutations in the *fc2-1* background.
Molecular and physiological phenotypes of the *fc2-1 atg* mutants were assessed. **A)** RT-qPCR analysis of transcripts from four-day-old seedlings grown under 6h light/18h dark light cycling conditions. Shown are mean values +/- SEM (n = 3 biological replicates). Closed circles represent individual data points. Statistical tests were not performed. **B)** Images of representative 35 and 42 day old plants grown in 24h constant light conditions.

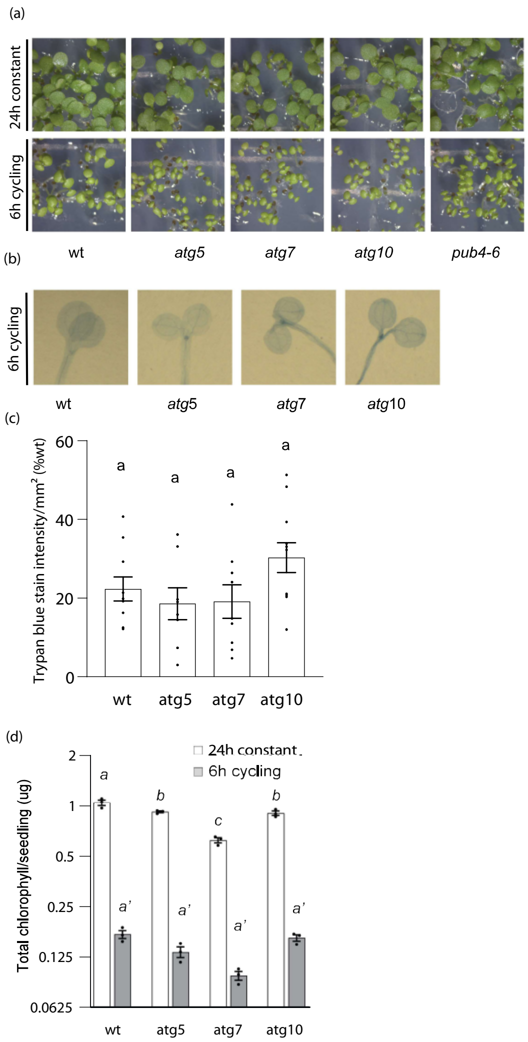


<u>Figure S2. Phenotypes of *atg5*, *atg7*, and *atg10* single mutant seedlings.</u>

The phenotypes of *atg* single mutant seedling were assessed in (24h) constant light and 6h light/18h dark (6h) cycling light. **A)** Seven-day old seedlings grown in 24h or 6h light conditions. **B)** Representative trypan blue stains of seedlings from panel A. The lack of dark blue color is indicative of healthy and alive cells. **C)** Mean values (+/- SEM) of the trypan blue signal in panel B ($n \geq 10$ seedlings). **D)** Mean total chlorophyll content (+/- SEM) of six-day old seedlings grown in 24h light or 6h cycling light conditions (n = 3 biological replicates). Statistical analyses were performed by one-way AVOVA tests followed by Tukey's HSD. Different letters above bars indicate significant differences (p value ≤ 0.05). For panel D, separate statistical analyses were performed for the different light treatments and the significance for the 6h cycling values is denoted by letters with a ′. In all bar graphs, closed circles represent individual data points.

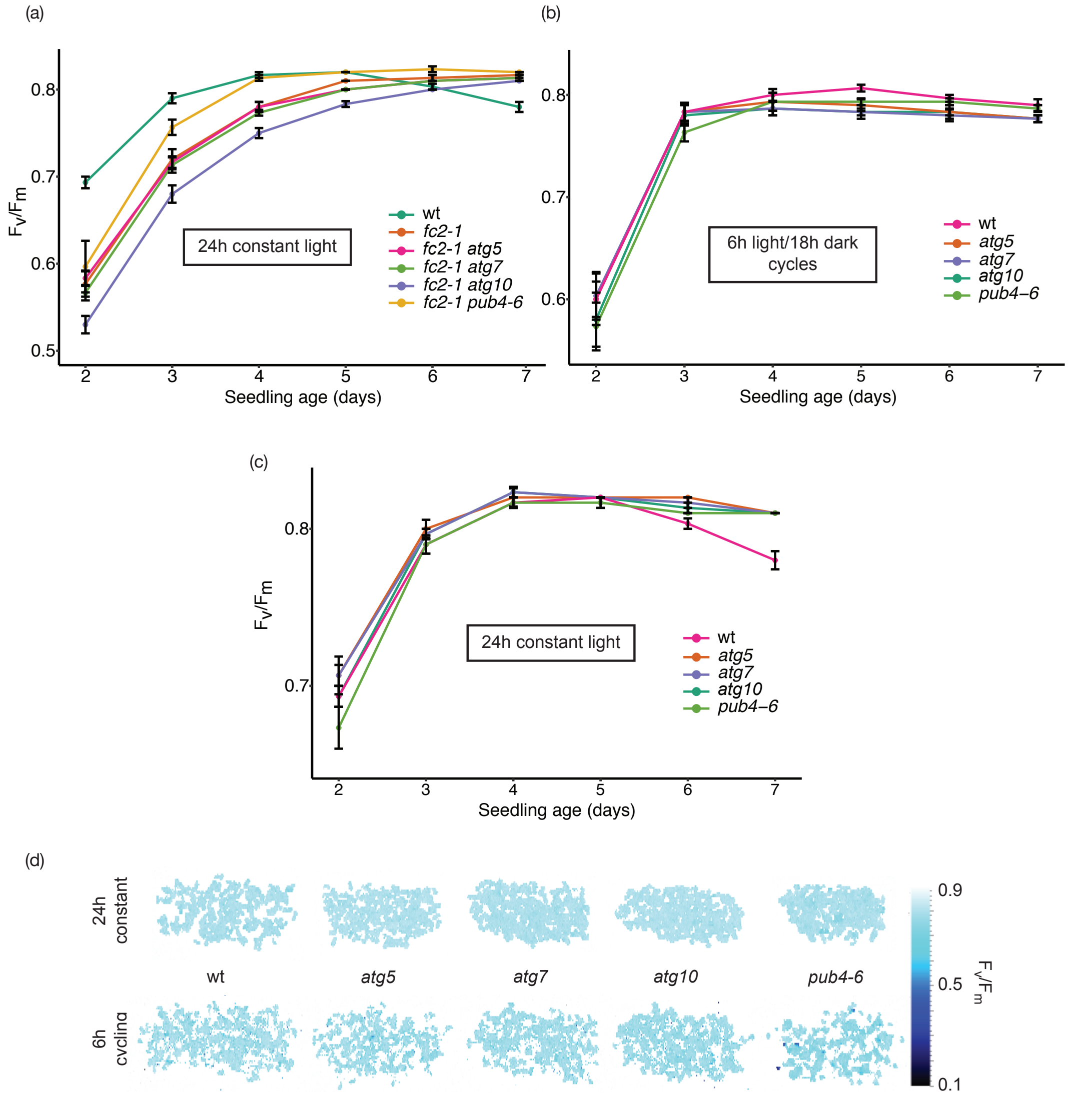


<u>Figure S3. Assessment of photosynthetic efficiency in the *atg* single and *fc2 atg* double mutants.</u> Maximum quantum yield of PSII ($F_v/F_m$) was measured in two to seven day old seedlings. **A)** Shown are the $F_v/F_m$ values of seedlings in the *fc2-1* background grown under 24h constant light conditions and **B)** seedlings in the wt background grown under 6h light/18h dark cycling or **C)** constant light conditions. Shown are mean values (+/- SEM) of biological replicates (n = 3). **D)** Representative images of maximum quantum yield of PSII ($F_v/F_m$) measured from three-day-old seedlings grown in the indicated light regiment.

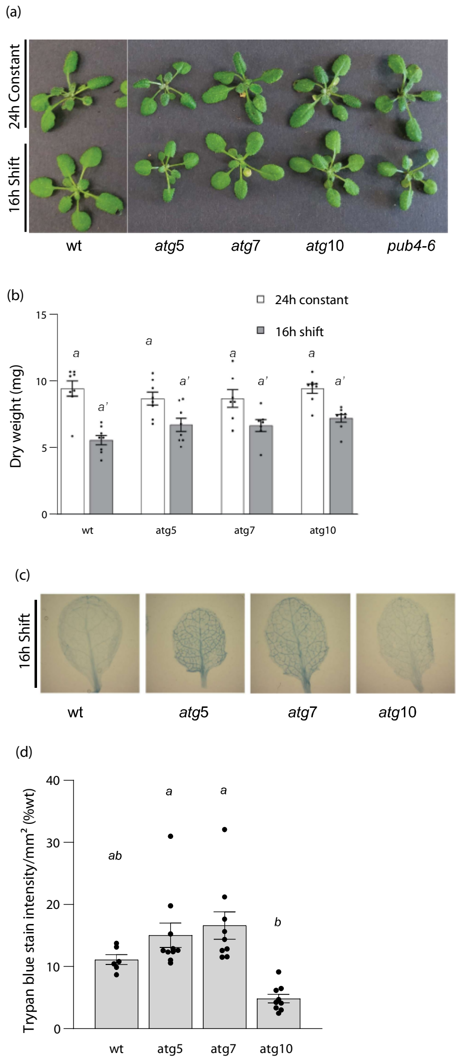


<u>Figure S4. Phenotypes of *atg5*, *atg7*, and *atg10* single mutant adult plants.</u>

The phenotypes of *atg* single mutant plants was assessed in the adult stage. **A)** Three-week old plants grown in 24h constant light or under stressed conditions (two weeks in 24h constant light and one week in 16h light/8h dark cycling light conditions). **B)** Mean biomass (+/- SD) of same plants (n = 8 plants). **C)** Representative trypan blue stains of single leaves from same plants. Lack of dark blue color is indicative of alive and healthy cells. **D)** Quantification of mean trypan blue signal (+/- SEM) in panel C (n $\geq$ 6 leaves from individual plants). Statistical analyses were performed by one-way AVOVA tests followed by Tukey's HSD. Different letters above bars indicate significant differences (p value $\leq$ 0.05). For panel B, separate statistical analyses were performed for the different light treatments and the significance for the light stressed group is denoted by letters with a ′. In all bar graphs, closed circles represent individual data points.

Table S1. List of autophagy-related genes and expression data used in heatmap generation.

| | | | **Expression relative to wt (from [1])** | | |
|---|---|---|---|---|---|
| **locus** | **Gene name** | **description** | ***fc2-1* 0 min** | ***fc2-1* 30 min** | ***fc2-1* 120 min** |
| *AT1G04140* | *AT1G04140* | *WD-40 repeat family protein;(source:Araport11)* | 1.083 | 1.004 | 0.883 |
| *AT1G04300* | *MUSE13* | *TUMOR NECROSIS FACTOR RECEPTOR-ASSOCIATED FACTOR 1B* | 0.963 | 1.034 | 1.161 |
| *AT1G22740* | *RAB7* | *RAB GTPASE 7* | 1.083 | 0.959 | 1.494 |
| *AT1G50030* | *TOR* | *TARGET OF RAPAMYCIN* | 1.028 | 0.996 | 1.093 |
| *AT1G54210* | *ATG12A* | *AUTOPHAGY 12A* | 0.962 | 0.884 | 1.132 |
| *AT1G60490* | *VPS34* | *VACUOLAR PROTEIN SORTING 34* | 1.028 | 1.027 | 1.069 |
| *AT1G73190* | *TIP3* | *ALPHA-TONOPLAST INTRINSIC PROTEIN 3* | 0.367 | 3.591 | 0.886 |
| *AT1G77890* | *ATG14A* | *AUTOPHAGY 14A* | 1.038 | 1.011 | 1.205 |
| *AT2G31260* | *ATG9* | *AUTOPHAGY 9* | 1.130 | 0.697 | 1.176 |
| *AT2G38470* | *WRKY33* | *WRKY DNA-BINDING PROTEIN 33* | 0.923 | 0.942 | 2.082 |
| *AT2G39780* | *RNS2* | *RIBONUCLEASE 2* | 1.000 | 1.087 | 0.895 |
| *AT2G40810* | *ATG18C* | *AUTOPHAGY 18C (YEAST HOMOLOG)* | 0.953 | 0.925 | 1.048 |
| *AT2G41980* | *SINAT1* | *SEVEN IN ABSENTIA OF ARABIDOPSIS THALIANA 1* | 1.072 | 0.782 | 0.899 |
| *AT2G44140* | *ATG4A* | *AUTOPHAGY 4A* | NA | NA | NA |
| *AT2G45170* | *ATG8E* | *AUTOPHAGY 8E* | 0.809 | 1.065 | 1.399 |
| *AT2G45260* | *COST1* | *CONSTITUTIVELY STRESSED 1* | 0.876 | 1.174 | 1.010 |
| *AT2G45980* | *ATI1* | *ATG8-INTERACTING PROTEIN 1* | 0.940 | 1.176 | 1.221 |
| *AT2G46240* | *BAG6* | *BCL-2-ASSOCIATED ATHANOGENE 6* | 1.015 | 0.444 | 11.649 |
| *AT3G01090* | *KIN10* | *SNF1-RELATED PROTEIN KINASE 10* | 0.917 | 0.987 | 1.098 |
| *AT3G06420* | *ATG8H* | *AUTOPHAGY 8H* | 0.857 | 1.108 | 1.386 |
| *AT3G07525* | *ATG10* | *AUTOPHAGY 10* | NA | NA | NA |
| *AT3G08850* | *RAPTOR1B* | *Regulatory-associated Protein of TOR 1B* | 1.006 | 0.928 | 1.058 |
| *AT3G09840* | *CDC48A* | *CELL DIVISION CYCLE 48A* | 0.946 | 1.133 | 0.981 |
| *AT3G12010* | *AT3G12010* | *Derives from AT3G12010; (source:Araport11)* | 1.058 | 0.889 | 0.933 |

| | | | | | |
|---|---|---|---|---|---|
| *AT3G13672* | *SINAT6* | *SEVEN IN ABSENTIA OF ARABIDOPSIS THALIANA 6* | 0.812 | 0.874 | 0.953 |
| *AT3G13970* | *ATG12B* | *AUTOPHAGY 12B* | NA | NA | NA |
| *AT3G15580* | *ATG8I* | *AUTOPHAGY 8I* | 0.865 | 1.071 | 1.362 |
| *AT3G18770* | *ATG13B* | *AUTOPHAGY 13B* | 0.940 | 1.138 | 0.880 |
| *AT3G19190* | *ATG2* | *AUTOPHAGY 2* | 1.129 | 0.927 | 1.232 |
| *AT3G21660* | *AT3G21660* | *UBX domain-containing protein;(source:Araport11)* | 0.967 | 1.187 | 0.859 |
| *AT3G28430* | *TT9* | *TRANSPARENT TESTA 9* | 0.994 | 0.967 | 0.835 |
| *AT3G49590* | *ATG13* | *AUTOPHAGY 13A* | 0.925 | 0.986 | 1.322 |
| *AT3G50590* | *TWD40-1* | *WD40/YVTN Repeat Protein* | 1.014 | 0.990 | 1.064 |
| *AT3G53230* | *CDC48B* | *CELL DIVISION CYCLE 48B* | 0.874 | 1.243 | 1.079 |
| *AT3G53930* | *ATG1B* | *AUTOPHAGY 1B* | 1.290 | 0.739 | 1.477 |
| *AT3G56440* | *ATG18D* | *AUTOPHAGY 18D (YEAST HOMOLOG)* | 1.168 | 0.964 | 1.009 |
| *AT3G57090* | *FIS1A* | *FISSION 1A* | 0.867 | 1.075 | 1.110 |
| *AT3G58040* | *SINAT2* | *SEVEN IN ABSENTIA OF ARABIDOPSIS THALIANA 2* | 1.125 | 0.817 | 1.071 |
| *AT3G59950* | *ATG4B* | *AUTOPHAGY 4B* | 0.975 | 0.969 | 1.035 |
| *AT3G60640* | *ATG8G* | *AUTOPHAGY 8G* | 0.939 | 1.266 | 1.257 |
| *AT3G61710* | *ATG6* | *AUTOPHAGY 6* | 1.092 | 0.872 | 1.054 |
| *AT3G61960* | *ATG1A* | *AUTOPHAGY 1A* | 1.090 | 0.872 | 1.014 |
| *AT3G62770* | *ATG18A* | *AUTOPHAGY 18A* | 0.983 | 1.128 | 1.223 |
| *AT4G02030* | *MTV16* | *MODIFIED TRANSPORT TO THE VACUOLE 16* | 1.047 | 0.754 | 1.129 |
| *AT4G04210* | *PUX4* | *PLANT UBX DOMAIN-CONTAINING PROTEIN 4* | 0.887 | 1.254 | 0.969 |
| *AT4G04620* | *ATG8B* | *AUTOPHAGY 8B* | 0.769 | 1.191 | 1.124 |
| *AT4G06676* | *AT4G06676* | *etoposide-induced protein;(source:Araport11)* | NA | NA | NA |
| *AT4G08540* | *ATG14B* | *AUTOPHAGY 14B* | NA | NA | NA |
| *AT4G15410* | *PUX5* | *PLANT UBX DOMAIN-CONTAINING PROTEIN 5* | 1.032 | 1.125 | 1.393 |
| *AT4G16520* | *ATG8F* | *AUTOPHAGY 8F* | 0.884 | 1.005 | 1.332 |
| *AT4G21980* | *ATG8A* | *AUTOPHAGY 8A* | 0.940 | 0.976 | 1.260 |
| *AT4G22150* | *PUX3* | *PLANT UBX DOMAIN-CONTAINING PROTEIN 3* | 0.951 | 1.047 | 1.171 |
| *AT4G22330* | *CES1* | *CERAMIDASE 1* | 0.975 | 0.936 | 1.153 |

| | | | | | |
|---|---|---|---|---|---|
| *AT4G24960* | *HVA22E* | *Homolog of ABA- and stress-inducible gene first isolated from barley.* | 0.898 | 1.204 | 0.715 |
| *AT4G29380* | *MTV11* | *MODIFIED TRANSPORT TO THE VACUOLE 11* | 0.977 | 1.153 | 1.173 |
| *AT4G30510* | *ATG18B* | *AUTOPHAGY 18B* | NA | NA | NA |
| *AT4G30790* | *ATG11* | *AUTOPHAGY 11* | 0.936 | 0.841 | 1.030 |
| *AT5G01770* | *RAPTOR1* | *Regulatory-associated Protein of TOR 1A* | 1.069 | 0.907 | 0.888 |
| *AT5G03340* | *CDC48C* | *CELL DIVISION CYCLE 48C* | 0.989 | 1.151 | 1.427 |
| *AT5G05150* | *ATG18E* | *AUTOPHAGY 18E (YEAST HOMOLOG)* | 0.876 | 1.069 | 0.716 |
| *AT5G12390* | *FIS1B* | *FISSION 1B* | 0.849 | 1.143 | 1.304 |
| *AT5G16280* | *TRS85* | *Tetratricopeptide Repeat (TPR)-like Superfamily Protein* | 0.916 | 1.114 | 1.246 |
| *AT5G17290* | *ATG5* | *AUTOPHAGY 5* | 1.106 | 0.876 | 1.257 |
| *AT5G24360* | *IRE1* | *INOSITOL REQUIRING 1-1* | 0.942 | 1.154 | 1.172 |
| *AT5G43560* | *MUSE14* | *TUMOR NECROSIS FACTOR RECEPTOR-ASSOCIATED FACTOR 1A* | 0.993 | 1.106 | 1.003 |
| *AT5G43930* | *RTP5* | *RESISTANT TO PHYTOPHTHORA 5* | 1.063 | 0.828 | 1.370 |
| *AT5G45900* | *ATG7* | *AUTOPHAGY 7* | 1.097 | 0.920 | 1.035 |
| *AT5G49540* | *AT5G49540* | *Rab5-interacting family protein;(source:Araport11)* | 0.920 | 1.099 | 1.057 |
| *AT5G50230* | *ATG16* | *AUTOPHAGY 16* | 0.927 | 1.084 | 1.077 |
| *AT5G61500* | *ATG3* | *AUTOPHAGY 3* | 1.018 | 0.976 | 1.076 |
| *AT5G66930* | *ATG101* | *AUTOPHAGY-RELATED 101* | 1.094 | 0.843 | 1.283 |

Table S2. List of predicted microautophagy-related genes and expression data used in heatmap generation.

| **Gene list derived from [2]** | | | **Expression relative to wt (from [1])** | | |
|---|---|---|---|---|---|
| **locus** | **gene name** | **description** | ***fc2-1*** **0 min** | ***fc2-1*** **30 min** | ***fc2-1*** **120 min** |
| *AT1G03950* | *VPS2.3* | *VACUOLAR PROTEIN SORTING-ASSOCIATED PROTEIN 2.3* | 0.984 | 0.874 | 1.184 |
| *AT1G07230* | *NPC1* | *NON-SPECIFIC PHOSPHOLIPASE C1* | 0.978 | 1.020 | 1.239 |
| *AT1G08190* | *VPS41* | *VACUOLAR PROTEIN SORTING 41* | 1.076 | 0.864 | 0.977 |
| *AT1G12470* | *VPS18* | *VACUOLAR PROTEIN SORTING 18* | 0.997 | 0.986 | 1.214 |

| | | | | | |
|---|---|---|---|---|---|
| *AT1G17730* | *VPS46.1* | *VACUOLAR PROTEIN SORTING 46.1* | 0.979 | 0.921 | 0.972 |
| *AT1G20110* | *FREE1* | *FYVE DOMAIN PROTEIN REQUIRED FOR ENDOSOMAL SORTING 1* | 1.002 | 0.969 | 1.206 |
| *AT1G54210* | *ATG12A* | *AUTOPHAGY 12A* | 0.962 | 0.884 | 1.132 |
| *AT1G60490* | *VPS34* | *VACUOLAR PROTEIN SORTING 34* | 1.028 | 1.027 | 1.069 |
| *AT1G73030* | *VPS46.2* | *VACUOLAR PROTEIN SORTING 46.2* | 0.898 | 1.043 | 1.134 |
| *AT1G77890* | *ATG14A* | *AUTOPHAGY 14A* | 1.038 | 1.011 | 1.205 |
| *AT2G06530* | *VPS2.3* | *SNF7 family protein;(source:Araport11)* | 1.026 | 0.976 | 1.147 |
| *AT2G19830* | *VPS32* | *SNF7 family protein;(source:Araport11)* | 1.024 | 0.875 | 1.209 |
| *AT2G27600* | *VPS4* | *VACUOLAR PROTEIN SORTING 4* | 1.078 | 0.881 | 1.136 |
| *AT2G31260* | *ATG9* | *AUTOPHAGY 12A* | 1.130 | 0.697 | 1.176 |
| *AT2G36680* | *AT2G36680* | *Modifier of rudimentary (Mod(r)) protein;(source:Araport11)* | 0.909 | 0.871 | 1.261 |
| *AT2G40810* | *ATG18C* | *AUTOPHAGY 18C* | 0.953 | 0.925 | 1.048 |
| *AT2G45170* | *ATG8E* | *AUTOPHAGY 8E* | 0.809 | 1.065 | 1.399 |
| *AT3G06420* | *ATG8H* | *AUTOPHAGY 8H* | 0.857 | 1.108 | 1.386 |
| *AT3G07525* | *ATG10* | *AUTOPHAGY 10* | NA | NA | NA |
| *AT3G08530* | *CHC2* | *CLATHRIN HEAVY CHAIN 2* | 1.029 | 1.032 | 1.029 |
| *AT3G09560* | *PAH1* | *PHOSPHATIDIC ACID PHOSPHOHYDROLASE 1* | 1.049 | 0.873 | 0.953 |
| *AT3G11130* | *CHC1* | *CLATHRIN HEAVY CHAIN 2* | NA | NA | NA |
| *AT3G12400* | *ELC* | *Ubiquitin-conjugating enzyme/RWD-like protein* | 0.928 | 1.043 | 1.149 |
| *AT3G15580* | *ATG8I* | *AUTOPHAGY 8I* | 0.865 | 1.071 | 1.362 |
| *AT3G18770* | *ATG13B* | *AUTOPHAGY 13B* | 0.940 | 1.138 | 0.880 |
| *AT3G19190* | *ATG2* | *AUTOPHAGY 2* | 1.129 | 0.927 | 1.232 |
| *AT3G45000* | *VPS24.2* | *SNF7 family protein;(source:Araport11)* | 1.033 | 1.000 | 0.973 |
| *AT3G49590* | *ATG13* | *AUTOPHAGY 13* | 0.925 | 0.986 | 1.322 |
| *AT3G53120* | *VPS28-2* | *Modifier of rudimentary (Mod(r)) protein;(source:Araport11)* | NA | NA | NA |
| *AT3G53930* | *ATG1B* | *AUTOPHAGY 1B* | 1.290 | 0.739 | 1.477 |
| *AT3G54860* | *VPS33* | *Sec1/munc18-like (SM) proteins superfamily* | 1.075 | 1.074 | 1.228 |
| *AT3G56440* | *ATG18D* | *AUTOPHAGY 18D* | 1.168 | 0.964 | 1.009 |

| | | | | | |
|---|---|---|---|---|---|
| *AT3G59950* | *ATG4B* | *AUTOPHAGY 4B* | 0.975 | 0.969 | 1.035 |
| *AT3G60640* | *ATG8G* | *AUTOPHAGY 8G* | 0.939 | 1.266 | 1.257 |
| *AT3G61710* | *ATG6* | *AUTOPHAGY 6* | 1.092 | 0.872 | 1.054 |
| *AT3G61960* | *ATG1A* | *AUTOPHAGY 1A* | 1.090 | 0.872 | 1.014 |
| *AT3G62080* | *CHMP7* | *CHARGED MULTI-VESICULAR BODY PROTEIN 7* | 1.067 | 0.767 | 1.435 |
| *AT3G62770* | *ATG18A* | *AUTOPHAGY 18A* | 0.983 | 1.128 | 1.223 |
| *AT4G04620* | *ATG8B* | *AUTOPHAGY 8B* | 0.769 | 1.191 | 1.124 |
| *AT4G05000* | *VPS28-2* | *Vacuolar protein sorting-associated protein VPS28 family protein* | 0.898 | 1.092 | 1.209 |
| *AT4G16520* | *ATG8F* | *AUTOPHAGY 8F* | 0.884 | 1.005 | 1.332 |
| *AT4G19003* | *VPS25* | *E2F/DP family winged-helix DNA-binding domain-containing protein;(source:Araport11)* | NA | NA | NA |
| *AT4G21560* | *VPS28-1* | *VACUOLAR PROTEIN SORTING 28* | 0.957 | 0.993 | 1.040 |
| *AT4G21980* | *ATG8A* | *AUTOPHAGY 8A* | 0.940 | 0.976 | 1.260 |
| *AT4G24690* | *NBR1* | *NEXT TO BRCA1 GENE 1* | 0.953 | 1.106 | 1.132 |
| *AT4G27040* | *VPS22* | *EAP30/Vps36 family protein;(source:Araport11)* | 0.917 | 1.042 | 0.997 |
| *AT4G29160* | *SNF7.1* | *SNF7 family protein;(source:Araport11)* | 0.963 | 1.006 | 1.198 |
| *AT4G29380* | *VPS15* | *WD-40 repeat family protein;(source:Araport11)* | 0.977 | 1.153 | 1.173 |
| *AT4G30790* | *ATG11* | *AUTOPHAGY 11* | 0.936 | 0.841 | 1.030 |
| *AT4G36630* | *VPS39* | *VACUOLAR PROTEIN SORTING 39* | 0.995 | 0.851 | 1.178 |
| *AT5G02500* | *HSC70* | *HEAT SHOCK COGNATE PROTEIN 70-1* | 1.087 | 1.046 | 1.031 |
| *AT5G04920* | *VPS36* | *EAP30/Vps36 family protein;(source:Araport11)* | 1.007 | 0.919 | 0.967 |
| *AT5G05150* | *ATG18E* | *AUTOPHAGY 18E* | 0.876 | 1.069 | 0.716 |
| *AT5G09260* | *VPS20.2* | *VACUOLAR PROTEIN SORTING-ASSOCIATED PROTEIN 20.2* | NA | NA | NA |
| *AT5G13860* | *VSP23B* | *ELCH-like protein;(source:Araport11)* | NA | NA | NA |
| *AT5G17290* | *ATG5* | *AUTOPHAGY 5* | 1.106 | 0.876 | 1.257 |
| *AT5G22950* | *VPS24.1* | *SNF7 family protein;(source:Araport11)* | 0.976 | 0.838 | 1.036 |

| | | | | | |
|---|---|---|---|---|---|
| *AT5G42870* | *PAH2* | *PHOSPHATIDIC ACID PHOSPHOHYDROLASE 2* | 1.188 | 0.781 | 1.451 |
| *AT5G44560* | *VPS2.2* | *SNF7 family protein;(source:Araport11)* | 1.019 | 0.940 | 1.129 |
| *AT5G45900* | *ATG7* | *AUTOPHAGY 7* | 1.097 | 0.920 | 1.035 |
| *AT5G46860* | *VAM3* | *Syntaxin/t-SNARE family protein* | 0.993 | 0.976 | 1.295 |
| *AT5G50230* | *ATG16* | *AUTOPHAGY 16* | 0.927 | 1.084 | 1.077 |
| *AT5G53330* | *Ub-EF1B* | *Ubiquitin-associated/translation elongation factor EF1B protein;(source:Araport11)* | 0.930 | 0.883 | 1.237 |
| *AT5G61500* | *ATG3* | *AUTOPHAGY 3* | 1.018 | 0.976 | 1.076 |
| *AT5G63880* | *VPS20.1* | *SNF7 family protein;(source:Araport11)* | 0.980 | 0.969 | 1.263 |
| *AT5G66930* | *ATG101* | *AUTOPHAGY-RELATED 101* | 1.094 | 0.843 | 1.283 |

Table S3. Mutant lines used in this study

| **mutant** | **Gene/locus** | **Mutation** | **Effect of mutation** | **reference** |
|---|---|---|---|---|
| *fc2-1* | *FC2/* AT2G30390 | GABI_766H08 T-DNA in 5'UTR | Reduced expression of *FC2* transcript | [3] |
| *pub4-6* | *PUB4/* AT2G23140 | c9847535t | G225R | [1] |
| *toc33* | *TOC33/PPI1/* AT1G02280 | c449839t | splice change, 3' end 2nd intron | [1] |
| *atg5-1* | *ATG5/* AT5G17290 | SAIL_129_B07 T-DNA intron 4 of 8 | Loss of transcript | [4] |
| *atg7-2* | *ATG7/* AT5G45900 | GABI_655B06 T-DNA exon 7 of 11 | Loss of transcript | [5] |
| *atg-1* | *ATG10/* AT3G07525 | SALK_084434 T-DNA exon 5 of 6 | Loss of transcript | [6] |

Table S4. Primers used in study

| **Gene** | **Oligo** | **Sequence** |
|---|---|---|
| **Genotyping Primers** | | |
| Salk LB | JP31 / LBb1.3 | ATTTTGCCGATTTCGGAAC |
| Sail LB | JP848 / LB | GCCTTTTCAGAAATGGATAAATAGCCTTGCTTCC |
| GABI-KAT LB | JP286 LB 08409 | ATATTGACCATCATACTCATTGC |
| GABI-KAT RB | JP285 RB 03144 | GTGGATTGATGTGATATCTCC |

| | | |
|---|---|---|
| *pub4-6* | LP JP | TATTAGAGTAGTGTGAGTCAGG |
| | RP JP | GATCCAGTGATTGTGTCATCC (*Hpa*II restriction enzyme cuts wt sequence) |
| *atg5-1* SAIL_129_B07 | LP JP1172 | GGAGCTTAACAAAGGGAAACG |
| | RP JP1173 | ACATAACCAATCGTTCCCTCC |
| *atg7-2* GABI_655B06 | LP JP1168 | GCCTTTTCAGAAATGGATAAATAGCCTTGCTTCC |
| | RP JP1169 | GGAGCTTAACAAAGGGAAACG |
| *atg10-1* SALK_084434 | LP JP1170 | ACATAACCAATCGTTCCCTCC |
| | RP JP1171 | CGTGTAACAGTGCATTGTTGG |
| **qPCR primers** | | |
| *ACTIN2* *AT3G18780* | For JP199 | GCACTTGCACCAAGCAGCAT |
| | Rev JP200 | CCTTTCAGGTGGTGCAACGAC |
| **Stress Response Markers** | | |
| *SIB1* *AT3G56710* | For JP589 | CAACCGGAGCCCATCTATT |
| | Rev JP590 | GGAGAAAGGTTGTGGTCGTC |
| *HSP26.5* *AT1G52560* | For JP585 | CGAGCTTATCGTTGCCTGAT |
| | Rev JP586 | CTCCGCCTTAATGTCCTCAA |
| *BAP1* *AT3G61190* | For JP338 | ATTGATGGATACGGTGGCCG |
| | Rev JP339 | CAGACCCCAAACCGGAACTC |
| *ATPase* *AT3G28580* | For JP336 | GAAGATCGGAAAAGCGTGGAA |
| | Rev JP337 | CCGGGTGGTCCAAACAAAAG |
| *ZAT12* *AT5G59820* | For JP344 | GCGTTGGTTACACGCGCTT |
| | Rev JP345 | CTTCAACGTAGTCACCGTGGG |
| *CYC8* *AT4G37370* | For JP1130 | AATGGGCATTGTCGAACGTG |
| | Rev JP1131 | TGGTCGAAATGCCAAACTCC |
| **Starvation Response Markers** | | |
| *C-starve* | | |
| *DIN6* *AT3G47340* | For WLO1757 | GAGTTCCACTTCTCGGTGCA |
| | Rev WLO1758 | GTGAGGGAAGATATGCCCCG |
| *WCOR413* *AT4G37220* | For WLO1759 | AAGGGGTGAGTTTTTGGCCA |
| | Rev WLO1760 | CCAAACCGGGTAAACGAGGA |

| | | |
|---|---|---|
| *DRM1*<br>*AT1G28330* | For WLO1761 | GGGATGATGTTGTGGCTGGA |
| | Rev WLO1762 | TCACCGCTGTACAACCAGTC |
| DIN11<br>AT3G49620 | For WLO1826 | AAAACGTGGACGGTGATTGG |
| | Rev WLO1827 | ACATGTCACCGATGTTGCAG |
| DIN10<br>AT5G20250 | For WLO1828 | TCGCACCGTTAAGTTGCATC |
| | Rev WLO1829 | ATAAAACGCGTCCCAAGTGC |
| *N-starve* | | |
| NRT2.1<br>AT1G08090 | For WLO1838 | TTTTTGCCTGGCGACGTTTG |
| | Rev WLO1839 | ACACAATGGGCATGAGCAAC |
| NRT2.4<br>AT5G60770 | For WLO1840 | AATGGCCGATGGTTTTGGTG |
| | Rev WLO1841 | TTGGCTTTGTGTTCGGTGTC |
| NRT2.5<br>AT1G12940 | For WLO1842 | TTTTGCTGCTTCGTCTCCAC |
| | Rev WLO1843 | ACGATACGAGCGAAAACAGC |
| ***Autophagy-related Genes*** | | |
| *BAG6*<br>*AT2G46240* | For WLO1630 | GGACCCGGAGAATGCTAGTG |
| | Rev WLO1631 | TTGCACGACGTCTGATCTGT |
| *WRKY33*<br>*AT2G38470* | For WLO1632 | CCAAACCGAGACTCGTCCAA |
| | Rev WLO1633 | TGCACTACGATTCTCGGCTC |
| *RAB7*<br>*AT1G22740* | For WLO1634 | AATTCTTGGAGACAGCGGGG |
| | Rev WLO1635 | AACCTCCTCTTTGCTCAGGC |
| *ATG1B*<br>*AT3G53930* | For WLO1636 | TCTCAAGAAGACGGGTTGCC |
| | Rev WLO1637 | TCCAGAGACCCATGGGAAGT |
| *CDC48*<br>*AT5G03340* | For WLO1638 | GGCTCGTCAATCTGCTCCTT |
| | Rev WLO1639 | CTGCAGCGTCTGAGCAAAAG |
| ***Core Autophagy Genes*** | | |
| *ATG5*<br>*AT5G17290* | For WLO1751 | ACAAATCCGACGTCGCTTCT |
| | Rev WLO1752 | CTCCGTCTCAGGCACAACTT |
| *ATG7*<br>*AT5G45900* | For WLO1753 | TGCCTTCTTCTTGGAGCTGG |
| | Rev WLO1754 | CAACCACGTCGTTGCAGAAG |

| | | |
|---|---|---|
| *ATG10 AT3G07525* | For WLO1755 | TCGAGAGGTCAGCGATGGTA |
| | Rev WLO1756 | ATCCAGTCCTCAGTCCCACA |
| *ATG3 AT5G61500* | For WLO1787 | GATGGTTGGCTGGCTACACA |
| | Rev WLO1788 | GGTTCTACTCCACGCGACAT |
| *ATG4a AT2G44140* | For WLO1789 | TCGTTTCTGGCAGCGAAGAT |
| | Rev WLO1790 | ATCGCTGTGTGGGTTTGAGT |
| *ATG4b AT3G59950* | For WLO1791 | CGTGTATTGGCCGCATTCAG |
| | Rev WLO1792 | GTTCACTCTGTCAAGCCCGA |
| *ATG8a AT4G21980* | For WLO1793 | CCTCTCGAGGCAAGGATGAG |
| | Rev WLO1794 | TCAAGCAACGGTAAGAGATCCA |
| *ATG8f AT4G16520* | For WLO1795 | GAAGAGAAGGGCAGAGGCTG |
| | Rev WLO1796 | CCAAATGTGTTTTCTCCGCTGT |
| *ATG12a - AT1G54210* | For WLO1797 | GGAGTCGTCGTCCCCGAG |
| | Rev WLO1798 | TCATCAGGGTTTGGCGAGAA |
| *ATG12b - AT3G13970* | For WLO1799 | GGCGACCGAATCTCCGAATT |
| | Rev WLO1800 | CTGATTCATCCGGGTTTGGC |
| *ATG2 - AT3G19190* | For WLO1801 | CAAAGGGCCGGGATACAACT |
| | Rev WLO1802 | CAGGTGACGAGACCAGCTTT |
| ***Predicted Microautophagy-related Genes*** | | |
| *ELC AT3G12400* | For WLO1850 | TTCGGCTGATCAGTCATTGC |
| | Rev WLO1851 | TGGTGAACATGCTGCACTTG |
| *VPS25 AT4G19003* | For WLO1852 | AGCTTCAATGCTTGCATGCC |
| | Rev WLO1853 | AGAGGAACCTTGTTCCAAGTGG |
| *VPS20.1 AT5G63880* | For WLO1854 | AACCCAAAGACGCAAGCTTG |
| | Rev WLO1855 | TCTTGCAGCTTGCTTTTCCG |
| *NBR1 AT4G24690* | For WLO1856 | ACTGGCGCTCATTCAAAGAC |
| | Rev WLO1857 | AAACACGACGAGGATGCTTG |
| *VPS15 AT4G29380* | For WLO1858 | TGCGTCAATTGCTTCTGAGG |
| | Rev WLO1859 | TCTTTCGCTGTTTCGCCATG |

| | | |
|---|---|---|
| *NPC1 AT1G07230* | For WLO1860 | TGAGCACGGTGGGTTTTATG |
| | Rev WLO1861 | AACACCCAATCGGTCAAACC |
| *VAM3 AT5G46860* | For WLO1862 | AGATTCGACACAAGCCGTTG |
| | Rev WLO1863 | TGCAACCTTGTCTTGTGCAG |
